# Nuclear PHGDH promotes neutrophil recruitment to drive liver cancer progression

**DOI:** 10.1101/2021.10.17.464745

**Authors:** Hongwen Zhu, Hua Yu, Hu Zhou, Wencheng Zhu, Xiongjun Wang

**Affiliations:** Department of Analytical Chemistry and CAS Key Laboratory of Receptor Research, Shanghai Institute of Materia Medica, Chinese Academy of Sciences, 555 Zuchongzhi Road, Shanghai 201203, China; Precise Genome Engineering Center, School of Life Sciences, Guangzhou University, Guangzhou 510006, China; Institute of Neuroscience, State Key Laboratory of Neuroscience, CAS Center for Excellence in Brain Science and Intelligence Technology, Shanghai Institutes for Biological Sciences, Chinese Academy of Sciences, Shanghai 201203, China

**Author notes:** Address correspondence to: Xiongjun Wang, Wencheng Zhu or Hu Zhou. Xiongjun Wang, Precise Genome Engineering Center, School of Life Sciences, Guangzhou University, Guangzhou 510006, China. (XW). Wencheng Zhu, Institute of Neuroscience, CAS Center for Excellence in Brain Science and Intelligence Technology, Chinese Academy of Sciences, Shanghai 201203, China. (WZ). Department of Analytical Chemistry and CAS Key Laboratory of Receptor Research, Shanghai Institute of Materia Medica, Chinese Academy of Sciences, 555 Zuchongzhi Road, Shanghai 201203, China. (HZhou). **Authorship note:** HZ and HY contributed equally to this work.

**Keywords:** nucleus PHGDH, cMyc, Liver cancer, CXCL1/8, Neutrophil

## Abstract

Metabolic dysregulation and the communications between cancer and immune cells are emerging as two essential features of malignant tumors. In this study, we observed that nuclear localization of phosphoglycerate dehydrogenase (PHGDH) associates with poor prognosis of liver cancer patients, and Phgdh is required for liver cancer progression in a mouse model. Unexpectedly, the impairment of Phgdh enzyme activity exerts a slight effect on liver cancer model, indicating PHGDH contributes to liver cancer progression mainly depending on its non-metabolic roles with nuclear location. PHGDH uses its ACT domain to bind cMyc in nuclear and forms a transactivation axis “PHGDH/p300/cMyc/AF9” which drives *CXCL1/8* gene expression. Chemokines CXCL1/8 promotes neutrophils recruitment and then supports tumor associated macrophages (TAMs) filtration in liver, thereby urging liver cancer into advanced stages. Forced cytosolic location of PHGDH or destruction of the PHGDH/cMyc interaction abolishes the oncogenic function of nuclear PHGDH. Collectively, this study reveals a non-metabolic role of PHGDH with altered cellular location in liver cancer, and suggests a promising drug target for liver cancer therapy by targeting the interaction between PHGDH and “undruggable” cMyc.

## Introduction

Primary liver cancer, including hepatocellular carcinoma (HCC) and intrahepatic cholangiocarcinoma (ICC), is one of the major causes of cancer-related death worldwide ^1^. Chemotherapy drugs are also limited in availability and are usually less effective than surgery or transplantation ^2^. During the past decades, genetic studies have uncovered several signaling pathways involved in hepatocarcinogenesis, such as the Wnt/β-catenin, MET and YAP/Hippo pathways ^3–5^. A method combining hydrodynamic gene delivery and Sleeping Beauty (SB)-mediated somatic integration has been widely used for long-term gene expression in mouse hepatocytes in order to develop murine models for liver cancer research ^6^.

In recent years, the omics field has been largely driven by technological advances, including RNA-seq for transcriptomic analysis and mass spectrometry (MS) technique for rapidly advancing proteomic applications ^7^. Through the combined analysis of protemics and RNA profling, we observed two enzymes of the serine synthesis pathway (SSP), Phgdh and Psat1, are upregulated in the advanced stage of mouse liver cancer model. The SSP has been demonstrated as a metabolic vulnerability in EGFR-mutant cancer ^8^. PHGDH is the first rate-limiting enzyme of the SSP and plays an important role in tumor resistance to serine starvation ^9^, reactive oxygen species (ROS) imbalance ^10^, sorafenib resistance ^11^ and brain metastasis ^12^. However, whether and how Phgdh regulates the progression of liver cancer is elusive.

Metabolic dysregulation reshapes tumor cells to enhance their growth and viability ^13^, but whether metabolic enzymes contribute to tumorigenesis through nonmetabolic functions is still relatively unclear. Some metabolic enzymes act independently of their classic metabolic activity to regulate tumor growth, survival and metastasis ^14,15^.

In addition to hepatocytes, there are non-parenchymal cells in liver tissue, mainly including macrophages, monocytes, neutrophils, lipid storage cells and sinusoidal endothelial cells. Normally, the stroma is involved in maintaining tissue homeostasis; however, when hepatocytes start to become cancerous, the surrounding stroma changes in ways that support tumor development ^16,17^. All of these cancer cells, immune cells and stromal cells, including their secreted factors, forms the tumor microenvironment (TME) ^18^.

The communications between tumor cells and the surrounding TME are based on complex systemic networks. In addition to direct cell-to-cell contact, extracellular communications through secreted cytokines/chemokines plays a key role in reprogramming TME ^18^. Of note, in the TME, tumor cells have been shown to acquire the ability to produce growth-promoting chemokines and to express chemokine receptors. For example, melanoma cells has been found to express a number of chemokines, including CXCL1/2/3/8 and CCL2/5, which have been implicated in tumor growth and progression ^19^. In this study, liver cancer cells expressed and secreted CXCL1/2/8, IL1B and CCL5, most of which match those chemokines expressed by the melanoma cells.

We further proved that CXCL1/8 directed the recruitment of neutrophils and probably supported TAM filtration. We speculated that the reprogrammed TME by liver cancer cells facilitated liver cancer progressing into advanced stage. In our view, there is a complicated network between the tumor cells and immune cells by the interaction of chemokines-chemokines receptors. Once tumor cells encounter different stresses due to rapid growth, medicial treatment or other reasons, they would actively initiate and stimulate this network. In that case, the immune cells, such as macrophages, monocytes, neutrophils, or the stromal cells would be recruited in the tumor tissue and contribute to cancer progression.

Conclusively, in this study, different from the role of PHGDH in the cytoplasm, nuclear PHGDH mainly forms a complex with cMyc and drives gene expression which is required for recruiting neutrophils and TAMs, but does not contribute to Sorafenib resistance. Therefore, the liver cancer cells with nuclear PHGDH demonstrate a new way of how metabolic enzymes function in regulating the reciprocal action of tumor cells and microenvironment.

## Results

### Dynamic intermodulation of the SSP and the Cyto C family correlates with liver cancer progression

In order to explore the underlying mechanism of liver cancer progression, an integrative transcriptomic and proteomic analysis was performed to systematically elucidate the changes in C57BL/6 mouse liver tissues induced by human MET (hMET); a truncated β-catenin mutant, DN90-β-catenin (MET/CAT); and the SB transposase ^20^. The mice were sacrificed at week 0, 2 or 7 (W0, W2 or W7, respectively) after hydrodynamic injection, and then the liver tissues were collected for both RNA-seq and tandem mass tag (TMT) 10-plex-based quantitative proteomic analysis (Figure 1A). W2 and W7 were considered as an initial stage and a final stage of malignant transformation in liver, respectively. In total, 17,612 genes were identified by RNA-seq at all three time points, and 8,417 proteins were quantified from the proteomic data in all samples. The 7,751 overlapping genes between the transcriptomic and proteomic data were used for unbiased comparison (Supplemental Figure 1A). For proteomic data, the protein reporter intensities in each TMT labeling channel were adjusted by median normalization to correct for differences in sample loading (Supplemental Figure 1B). The correlation matrix and principal component analysis (PCA) both suggested good repeatability of sample quantification from the same time point and clear separation among different time points (Supplemental Figure 1, C and D). The correlation coefficients between the RNA and protein data were high: 0.63 in W2 and 0.39 in W7 (Supplemental Figure 1E).

**Figure 1.**
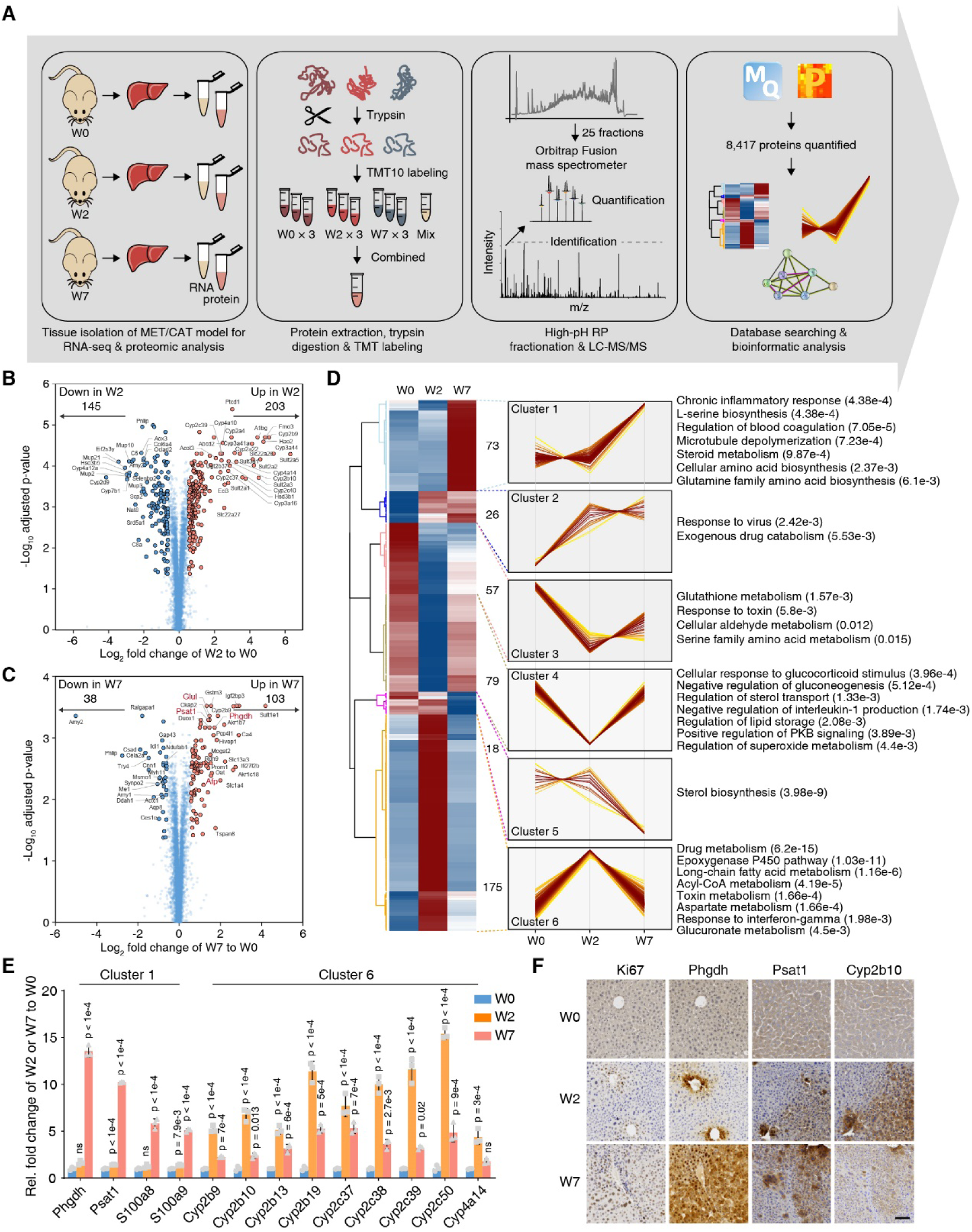
Dynamic intermodulation of the SSP and the Cyto C family correlates with liver cancer progression. **(A)** Experimental workflow for quantitative proteomic analysis. **(B)** The volcano plot presents the differentially regulated proteins in W2 vs. W0. The x-axis represents the log2-transformed ratio in W2 relative to W0. The y-axis represents the -log10-transformed p-value (adjusted by the Benjamini-Hochberg method). **(C)** The volcano plot presents the differentially regulated proteins in W7 vs. W0. **(D)** The differentially regulated proteins were separated into six clusters by HCA. Representative enriched biological processes for the proteins in each cluster are shown on the right. The p-value in the bracket was calculated by Fisher’s exact test. **(E)** Validation of the indicated mRNA changes by qRT-PCR in liver tissues of W0, W2 and W7 of the MET/CAT model. Rel., relative. (Mean ± SD, one-way ANOVA followed by Dunnett’s multiple comparisons test using W0 as the control, n=3.) **(F)** Immunohistochemistry (IHC) stainningof Ki67, Phgdh, Psat1 and Cyp2b10 in mouse liver sections of W0, W2 and W7 of the MET/CAT model. Scale bar, 200 μm.

Next, we analyzed the Kyoto Encyclopedia of Genes and Genomes (KEGG) pathways and Gene Ontology (GO) biological processes involved in the progression of MET/CAT-driven hepatocarcinogenesis. The gene expression in W2 reveals a more dramatic fluctuation than that in W7, both in RNA and protein levels (Supplemental Figure 1F). The most highly represented pathways and biological processes for the differentially expressed RNAs and proteins in W2 and W7 were liver detoxification-related pathways or biological processes, including chemical carcinogenesis, oxidation-reduction and the epoxygenase P450 pathway, and metabolism-related pathways, such as steroid hormone biosynthesis and retinol metabolism (Supplemental Figure 1G). The results of the combined transcriptomic and proteomic analysis led us to further focus on dysregulation of detoxification-related metabolism in the mechanism of liver cancer progression.

Proteins directly regulate biological behavior. Given the high degree of consistency between the RNA and protein data, we focused on the proteomic data to analyze the dynamic changes in proteins from W0 to W2 and then to W7 after SB induction. A total of 203 proteins and 145 proteins were up-regulated and down-regulated, respectively, in W2 compared to W0. As revealed in the volcano plot, members of the Cytochrome C family were significantly up-regulated in W2 (Figure 1B). In W7, 103 and 38 proteins were up- and down-regulated, respectively. The clinically relevant liver cancer biomarkers Afp and Glul were identified to be up-regulated in W7, suggesting the reliability of this liver cancer induction model. Notably, two key enzymes in L-serine (Ser) biosynthesis, Phgdh and Psat1, also showed obvious increases in W7 (Figure 1C). The differential protein expression analysis revealed that W2 and W7 represent two distinct states in liver cancer progression.

The differentially expressed proteins were further grouped into six clusters by hierarchical cluster analysis (HCA) (Figure 1D). Cluster 1 was a group of proteins that were markedly up-regulated in W7 and were enriched for biological processes such as the chronic inflammatory response, Ser biosynthesis, microtubule depolymerization, steroid metabolism and glutamine family amino acid biosynthesis. Interestingly, the Cluster 6 proteins were up-regulated in W2 but had returned to basal levels in W7 and were associated with drug metabolism, the epoxygenase P450 pathway, acyl-CoA metabolism, aspartate metabolism and the interferon-gamma response. Protein-protein interaction network analysis highlighted amino acid biosynthesis-related proteins, including Phgdh, Psat1, Oat and Glul, and inflammation-related proteins, such as S100a8 and S100a9 in Cluster 1 and members of the cytochrome C family in Cluster 6 (Supplemental Figure 1H). S100a8 and S100a9 have been reported to play protumorigenic roles in carcinogen-induced HCC ^21^. The expression changes in Phgdh, Psat1, S100a8, S100a9 and a few members of the cytochrome C family were validated by quantitative real-time PCR (qRT-PCR) (Figure 1E). Among these proteins, Ki67, Phgdh, Psat1 and Cyp2b10 were further confirmed to exhibit expression alterations by immunohistochemistry (IHC) (Figure 1F).

### Conditional deletion of Phgdh in hepatocytes attenuates MET/CAT-driven hepatocarcinogenesis, but inhibiting the enzyme activity of Phgdh does not efficiently prevent liver cancer progression

Phgdh and Psat1 are key enzymes, which execute the first two steps of serine (Ser) biosynthesis (Figure 2A) ^22^. To examine the functional roles of *Phgdh* in liver cancer, we induced liver tumorigenesis in *Phgdh*^*fl/fl*^ (control) and *Phgdh*^*LKO*^ mice by hydrodynamic injection. The *Phgdh*^*LKO*^ mice were constructed by deleting the second exon of *Phgdh* gene (Figure 2B). PCR, Western blot and IHC analyses confirmed the successful and specific knockout of *Phgdh* in mouse livers (Supplemental Figure 2, A-C). At the early time point (W2 after injection), the liver tissues of *Phgdh*^*LKO*^ mice showed a weakly reduced volume in the gross image. By the hematoxylin and eosin (H&E) staining, we observed some scattered tumor cells in the liver of *Phgdh*^*fl/fl*^ mouse (Figure 2C, left). Furthermore, compared to *Phgdh*^*LKO*^ mouse, we found a markedly increased liver volume and more tumor nodules in control mice (*Phgdh*^*fl/fl*^) at W7 after injection (Figure 2C, right). Loss of *Phgdh* in hepatocytes resulted in significant decreases in Ki67-positive cell numbers and liver/body weight ratios (Figure 2, D and E). In addition, *Phgdh* deletion significantly increased the ROS level in the liver (Figure 2F) ^10^. To verify the dysfunction of the SSP, we measured three metabolites, 3-phosphoglycerate (3PG), 3-phosphohydroxypyruvate (3PHP) and Ser in the SSP. Loss of *Phgdh* significantly blocked the SSP pathway, as evidenced by increased levels of 3PG and decreased levels of 3PHP and Ser (Figure 2G). As expected, deletion of *Phgdh* in hepatocytes prolonged survival time in mice after MET/CAT-driven hepatocarcinogenesis (Figure 2H). However, under normal living conditions, the liver/body weight ratios of 6-week-old *Phgdh*^*LKO*^ mice were comparable with controls (Supplemental Figure 2D). The H&E staining results (Supplemental Figure 2C) showed that the hepatocytes of the 6-week-old *Phgdh*^*LKO*^ mice was as normal as the controls. Furthermore, the serum ALT and AST levels in the livers of *Phgdh*^*fl/fl*^ and *Phgdh*^*LKO*^ mice did not differ (Supplemental Figure 2E), suggesting that liver functions of the *Phgdh*^*LKO*^ mice were apparently normal.

**Figure 2.**
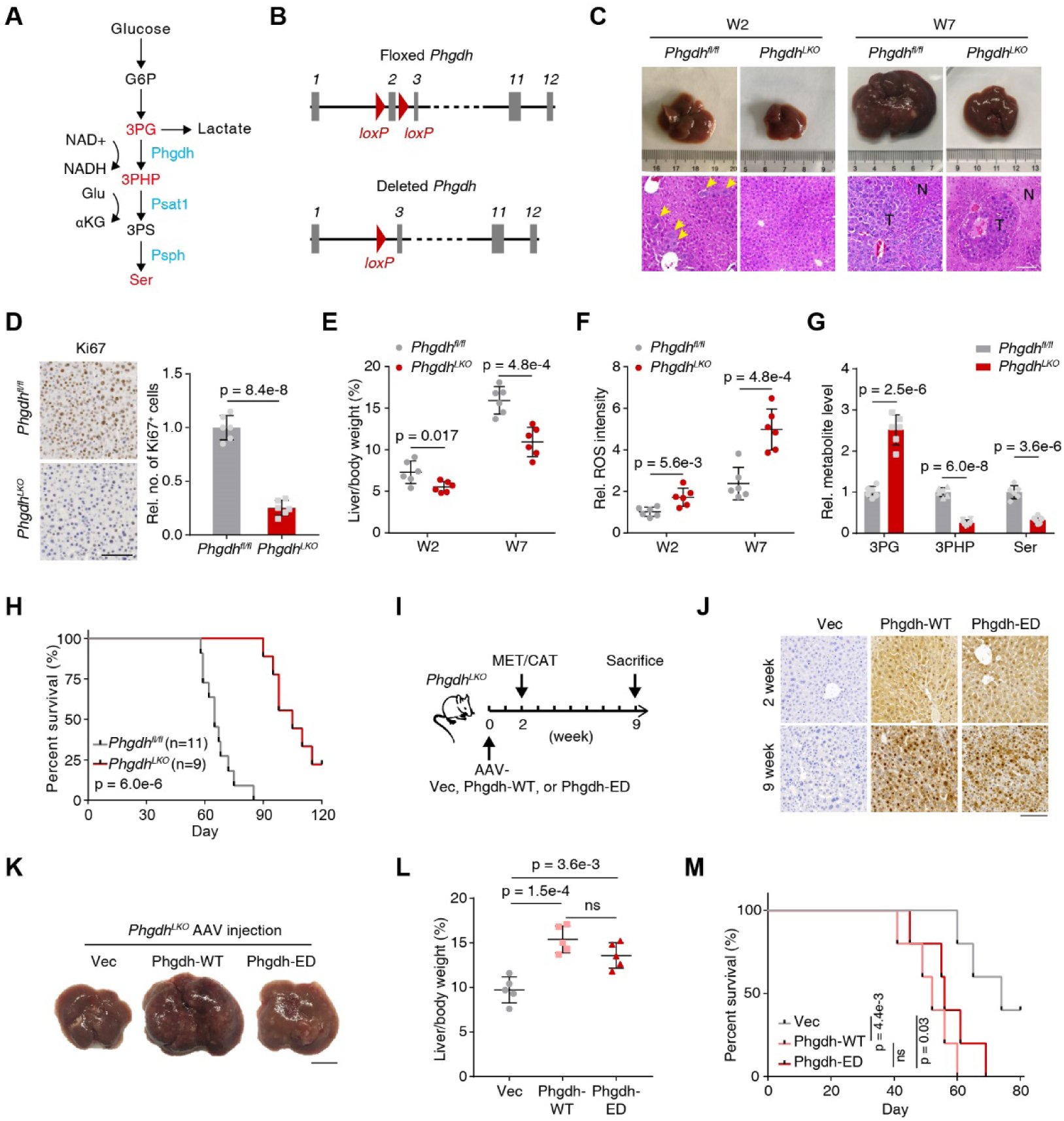
Inhibition of Phgdh activity does not efficiently hinder liver cancer progression induced by MET/CAT. **(A)** Metabolic enzymes Phgdh, Psat1 and Psph and the related metabolites in the SSP. G6P, glucose-6-phosphate; 3PS, 3-phosphoserine; nicotinamide adenine dinucleotide (oxidized form, NAD+; reduced form, NADH); Glu, glutamate; αKG, alpha-ketoglutarate. **(B)** Scheme of the *Phgdh* gene locus and related alleles. The *Phgdh* floxed alleles have two *loxP* sites (red triangles) flanking the second exon (gray boxes). Mice with *Phgdh* floxed alleles were crossed with a Cre line to generate the deleted *Phgdh* allele. **(C)** Macroscopic images and H&E staining of mouse (*Phgdh*^fl/fl^ and *Phgdh*^*LKO*^) liver sections at W2 and W7 after hydrodynamic injection of MET/CAT constructs and the SB transposase. Scale bar, 100 μm. **(D)** Ki67 staining of liver sections at week 7 after injection of MET/CAT (left panel). Relative numbers of Ki67-positive cells in liver sections (right panel). Scale bars: 200 μm. Rel., relative; no., number (Mean ± SD, two-tailed Student’s t-test, n = 6.) **(E)** Liver/body weight ratios of MET/CAT-transfected mice (*Phgdh*^fl/fl^ and *Phgdh*^*LKO*^) measured at W2 and W7. (Mean ± SD, two-tailed Student’s t-test, n = 6.) **(F)** The relative ROS intensities were measured in liver samples of *Phgdh*^fl/fl^ and *Phgdh*^*LKO*^ mice at W2 and W7 after injection of MET/CAT. (Mean ± SD, two-tailed Student’s t-test, n = 6.) **(G)** The relative levels of three metabolites (3PG, 3PHP and Ser) were measured in liver samples of *Phgdh*^fl/fl^ and *Phgdh*^*LKO*^ mice at W7 after injection of MET/CAT. (Mean ± SD, two-tailed Student’s t-test, n = 6.) **(H)** Kaplan-Meier plot showing the survival of *Phgdh*^fl/fl^ and *Phgdh*^*LKO*^ mice after injection of MET/CAT over 120 days. (Log-rank test) **(I)** Schematic diagram of AAV treatment in MET/CAT-induced liver tumor using *Phgdh*^*LKO*^ mice. The AAV treatment (Vec, Phgdh-WT and Phgdh-ED) was conducted in mice at W0. The mice were subjected to MET/CAT induction at W2 and sacrificed at W9. **(J)** IHC staining of Phgdh in mouse liver sections at W2 and W9 after *Phgdh*^*LKO*^ mice receving AAV treatments (Vec, Phgdh-WT and Phgdh-ED). Scale bar, 100 μm. **(K)** Macroscopic images of *Phgdh*^*LKO*^ mouse livers under different AAV treatments (Vec, Phgdh-WT and Phgdh-ED) at W9. Scale bar, 1 cm. **(L)** Liver/body weight ratios of MET/CAT-transfected mice under different AAV treatments (Vec, Phgdh-WT and Phgdh-ED) at W9 were measured. (Mean ± SD, one-way ANOVA followed by Tukey’s multiple comparisons test, n = 5) **(M)** Kaplan-Meier plot showing the survival of MET/CAT-transfected mice under different AAV treatments (Vec, Phgdh-WT and Phgdh-ED). (Log-rank test, n = 5)

To understand how Phgdh functions in MET/CAT-driven liver cancer progression, we administered NCT-503, a specific small-molecule inhibitor of Phgdh enzyme activity ^23^, to MET/CAT mice at W2 (Supplemental Figure 2F). The efficiency of Phgdh enzyme activity inhibition by NCT-503 was measured in mouse livers (Supplemental Figure 2G). Surprisingly, this inhibitor did not ultimately prevent the formation of liver cancer (Supplemental Figure 2H). Drug intervention at the early stage moderately delayed the tumor formation and only slightly decreased the tumor burden, as revealed by Ki67 staining and liver/body weight ratios (Supplemental Figure 2, I and J). Inhibition of Phgdh with NCT-503 did not promote the survival of mice bearing MET/CAT-induced liver cancer (Supplemental Figure 2K). Inhibiting the enzyme activity of Phgdh somewhat decreased liver ROS levels (Supplemental Figure 2L) and induced dysfunction of the SSP (Supplemental Figure 2M), suggesting that the compound worked *in vivo*. To specifically characterize the enzymatic activity of Phgdh in MET/CAT-induced tumorigenesis, we delivered adeno-associated viruses (AAV) carrying empty vector (Vec), wild-type Phgdh (Phgdh-WT) or enzyme-dead Phgdh (Phgdh-ED) under the regulation of the thyroid hormone binding globulin promoter into *Phgdh*^*LKO*^ adult mice (Figure 2I). AAV treatment specifically and efficiently drove the expression of Phgdh-WT or -ED in hepatocytes at week 2 (Figure 2J, upper). Then these mice were subjected to the MET/CAT to induce hepatocarcinogenesis (Figure 2I). In line with the data from inhibitor NCI-503, the restored Phgdh-WT in mouse liver has promoted the tumorigenesis, but the expression of Phgdh-ED also facilitated this process (Figure 2, K and L). Consequently, mice that received Phgdh-ED exhibited a markedly reduced lifespan compared to the controls (Figure 2M). Combined with data from NCI-503 treatment, the results suggest that there is another role of Phgdh involving in liver tumorigenesis independent of its enzymatic activity.

### PHGDH interacts with cMyc as a coactivator in a manner independent of its enzyme activity

In order to analyze the nonmetabolic effect of Phgdh, coimmunoprecipitation (Co-IP) with liquid chromatography (LC)-tandem MS (MS/MS) was performed to identify the proteins interacting with Phgdh during MET/CAT-induced liver cancer. cMyc is among the top enriched transcription factors driving HCC progression (Figure 3A) ^24–26^. Of note, in the Biological General Repository for Interaction Datasets (BioGRID) database (https://thebiogrid.org/), cMyc is also recorded as an interacting protein of Phgdh. We identified that cMyc interacted with Phgdh in the induced liver cancer group but not in the control group using a Co-IP assay (Figure 3B).

**Figure 3.**
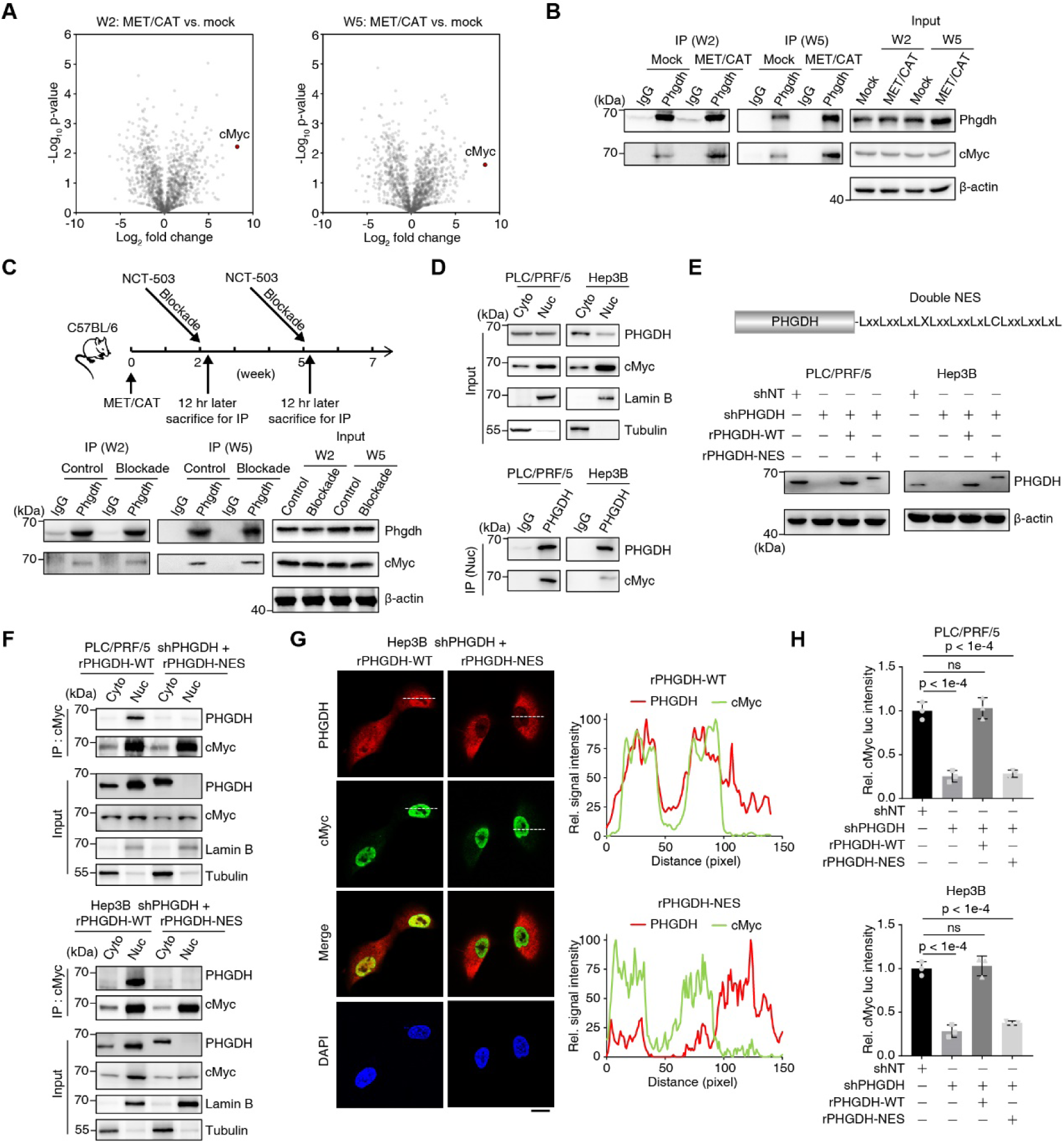
Phgdh interacts with cMyc in a manner independent of its enzyme activity. **(A)** Volcano plot of the proteins differentially interacting with Phgdh as determined by the ratio between the MET/CAT-induced liver cancer group and the mock group in W2 and W5. The mice in the mock group were injected with blank control plasmids. cMyc is shown as brown red circles. **(B)** Co-immunoprecipitation (Co-IP) assays were performed with an antibody against Phgdh followed by Western blot analysis of Phgdh to detect the interaction between Phgdh and cMyc. Loading control, β–actin. **(C)** C57BL/6 mice were injected with NCT-503 (dosage, 100 mg/kg) through the tail vein in W2 and W5, and 12 hours later after injection, mouse liver tissues were dissected for Co-IP assay to detect the interaction between Phgdh and cMyc. Loading control, β–actin. **(D)** Separation of nuclei and cytosol was performed using PLC/PRF/5 and Hep3B cells. Co-IP assay was conducted using nuclei to detect the interaction between Phgdh and cMyc. Cyto, cytosol; Nuc, nuclei. **(E)** The construction of nuclear export sequence (NES) tagged PHGDH (PHGDH-NES) in its C terminal was validated by immunoblotting in PLC/PRF/5 and Hep3B cells. shNT, a non-targeting short hairpin RNA (shRNA); shPHGDH, shRNA against PHGDH; rPHGDH-WT, shRNA-resistant wild type PHGDH; rPHGDH-NES, shRNA-resistant PHGDH-NES. Loading control, β–actin. **(F)** Separation of nuclei and cytosol was performed using PHGDH-depleted PLC/PRF/5 and Hep3B cells rescued with rPHGDH-WT or rPHGDH-NES. Co-IP assay was conducted using nuclei to detect the interaction between Phgdh and cMyc. **(G)** Immunofluorescence assay using PHGDH-depleted Hep3B cells rescued with rPHGDH-WT or rPHGDH-NES. Cells were fixed with 80% methanol and stained with antibodies against cMyc or PHGDH. 4′, 6-Diamidino-2-phenylindole (DAPI) was used as a nuclear localization marker (left panel). Scale bars: 20 μm. The fluorescence intensity profile of regions of interest (the dotted lines) was quantified to illustrate the colocalization of PHGDH and cMyc using ImageJ (right panel). **(H)** cMyc transactivation was measured with a Dual-Luciferase^®^ Reporter Assay System according to the manual using cells from **E**. (Mean ± SD, one-way ANOVA followed by Dunnett’s multiple comparisons test, n = 3)

Then, we used NCT-503 to block the enzyme activity of Phgdh in the liver and then harvested liver tissues for interaction analysis. The results showed that inhibiting the enzyme activity of Phgdh did not affect its binding to cMyc, suggesting that Phgdh has a nonmetabolic function (Figure 3C). Consistent with the results obtained in the induced mouse liver cancer model, PHGDH and cMyc interacted in the human liver cancer cell lines, and this interaction did not depend on the enzyme activity of PHGDH (Supplemental Figure 3A). We stably knocked down *PHGDH* in these cell lines (Supplemental Figure 3B), and a luciferase assay showed that the enzyme inactivation of PHGDH resulted in a decrease in the transactivation of cMyc by more than half (Supplemental Figure 3C).

We speculated that a prerequisite for the interaction of Phgdh with cMyc is its nuclear localization. Therefore, we extracted the nuclei of two human liver cancer cell lines (PLC/PRF/5, Hep3B) and found that PHGDH partially localized to the nuclei (Figure 3D). To export PHGDH from nuclei into cytosol, we artificially added double nuclear export sequence (NES) in the C terminal of PHGDH (Figure 3E). A Co-IP assay was performed using the separated fraction of cytosol and nuclei, which showed that the PHGDH-NES successfully located in the cytosol and the interaction between PHGDH-NES and cMyc was not observed (Figure 3F). The results of immunofluorescence analysis further showed that PHGDH and cMyc colocalized in the nucleus and PHGDH-NES mutant has no obivious co-localization (Figure 3G). Additionally, a luciferase assay showed that the exportation of PHGDH from nuclei to cytosol resulted in a decrease in the transactivation of cMyc (Figure 3H).

### The PHGDH ACT domain is required for forming the PHGDH/p300/cMyc/AF9 axis and regulates cMyc transactivation

Human PHGDH contains five domains: substrate-binding 1 (SB1), nucleotide-binding (NB), substrate-binding 2 (SB2), allosteric substrate binding (ASB), and aspartate kinase-chorismate mutase-tyrA prephenate dehydrogenase (ACT) (Figure 4A) ^27^. To clarify which region of PHGDH targeting cMyc, we separately deleted above five domains in the full-length of PHGDH. A Co-IP assay showed that the ACT domain containing 460–533 amino acids of PHGDH was required for its association with cMyc (Figure 4B).

**Figure 4.**
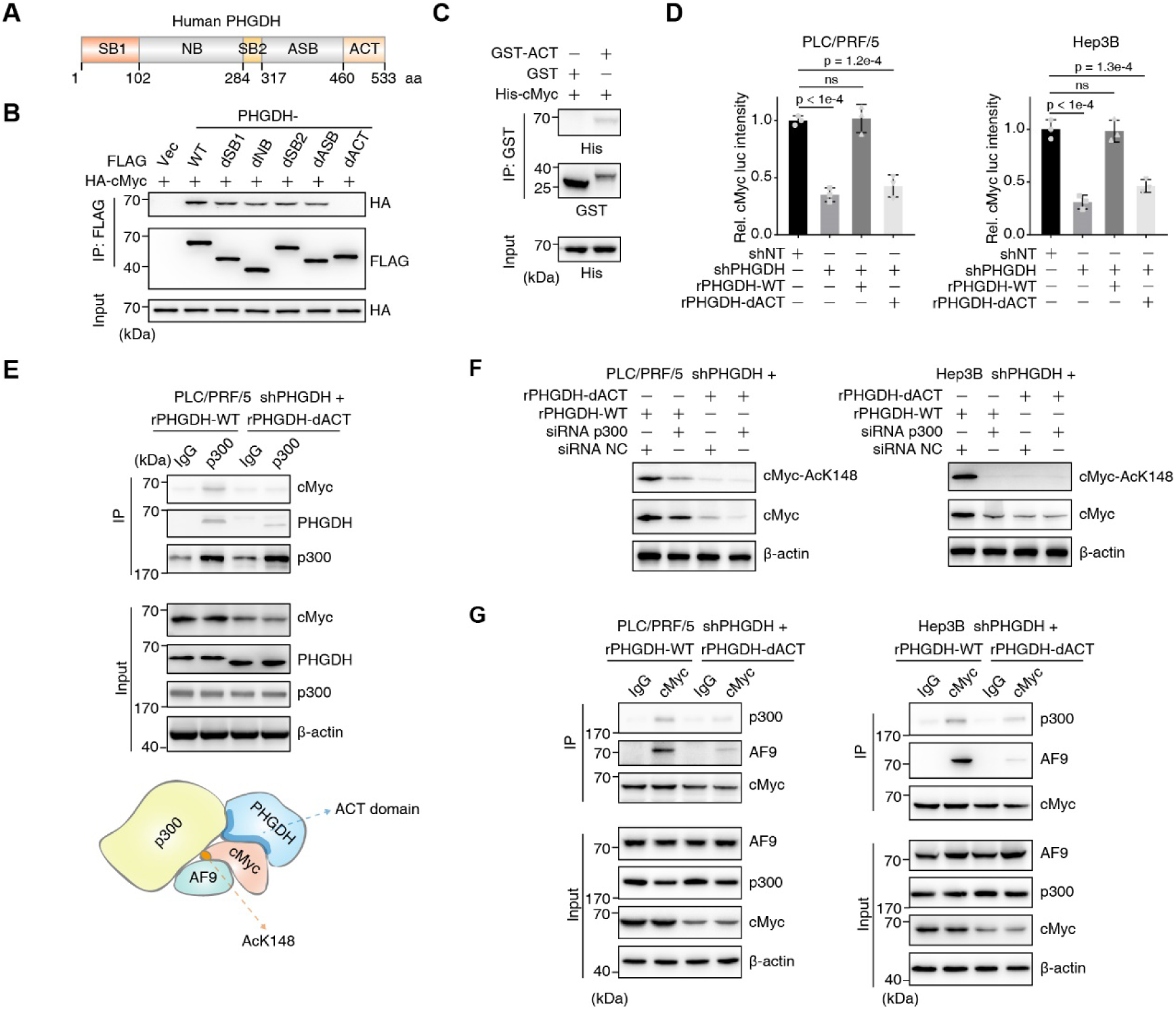
The PHGDH ACT domain is required for forming PHGDH/p300/cMyc/AF9 axis and regulates cMyc transactivation. **(A)** Schematic diagram of human PHGDH protein and its five domains. aa, amino acid. **(B)** HA-tagged cMyc and FLAG-tagged PHGDH (including WT and five domain truncates) were transiently transfected into HEK293T cells. Co-IP was performed with an antibody against FLAG. Antibodies against HA and FLAG were used to detect the association between cMyc and PHGDH. HA was used as an imput control. dSB1, SB1 domain depletion; dNB, NB domain depletion; dSB2, SB2 domain depletion; dASB, ASB domain depletion; dACT, ACT domain depletion. **(C)** GST-tagged ACT domain and His-tagged cMyc protein were purified from *E. coli*, and a GST pull-down assay was performed by incubating both the recombinant proteins together. GST was used as a blank control. Antibodies against His and GST were used to detect the association between cMyc and the ACT domain. **(D)** cMyc transcriptional activity was measured by the Dual-Luciferase^®^ Reporter Assay System according to the manual using PHGDH-depleted PLC/PRF/5 and Hep3B cells rescued with rPHGDH-WT or rPHGDH-dACT. Cells expressing shNT were used as control. rPHGDH-dACT, shRNA-resistant PHGDH-dACT. (Mean ± SD, one-way ANOVA followed by Dunnett’s multiple comparisons test, n = 3) **(E)** Co-IP analysis of p300, cMyc and PHGDH was performed using PHGDH-depleted PLC/PRF/5 cells rescued with rPHGDH-WT or rPHGDH-dACT. Antibody against p300 was used to enrich p300 associated complex. Immunoblotting analysis of PHGDH, cMyc and p300 was performed using the indicated antibodies. **(F)** p300 was transiently depleted by specific siRNA in PHGDH-depleted PLC/PRF/5 cells rescued with rPHGDH-WT or rPHGDH-dACT. Immunoblotting analysis of cMyc-AcK148, cMyc and p300 was performed using the indicated antibodies. NC, negative control. **(G)** Co-IP analysis of cMyc, p300 and PHGDH was performed using PHGDH-depleted PLC/PRF/5 cells rescued with rPHGDH-WT or rPHGDH-dACT. Antibody against cMyc was used to enrich cMyc associated complex. Immunoblotting analysis of PHGDH, p300, cMyc and AF9 was performed using the indicated antibodies.

The GST-tagged ACT domain was purified from *E. coli* and His-tagged cMyc protein was expressed in E. coli as inclusion bodies, solubilized, refolded, and further purified. A GST pull-down assay showed a significant interaction of cMyc with the PHGDH-ACT domain (Figure 4C). Importantly, the binding of the ACT domain to cMyc could be essential for PHGDH function in cMyc transactivation. We found that the capacity of PHGDH to transactivate cMyc was significantly diminished when we deleted PHGDH-ACT domain (Figure 4D). Meanwhile, cMyc protein levels in the cells expressing the mutant PHGDH-dACT were decreased (Supplemental Figure 4A). Given that the ACT domain promoted cMyc transactivation and cMyc recuits p300 to activate gene expression, we speculated that PHGDH ACT domain could associate with p300, but whether ACT domain integrates p300 and cMyc together is still unclear. To test this, a Co-IP assay was peformed and showed loss of ACT domain, p300 has no obvious interaction with cMyc and PHGDH (Figure 4E). As we observed a reduction of cMyc protein levels in the cells expressing PHGDH-dACT mutant, we further examined cMyc protein levels upon p300 depletion and a consistent result was shown, indicating PHGDH and p300 exert the effect on cMyc protein in the same axis (Figure 4F). We suspect that PHGDH uses its ACT domain to directly bind with cMyc in order to form a signaling axis. Additionally, our previous study reported that AF9 recruited p300 to acetylate cMyc at its K148 ^28^. Thus, we wondered whether p300/cMyc/AF9 axis associated with PHGDH. As shown in Figure 5G, comparing to PLC/PRF/5 cells expressing WT PHGDH, the cells expressing PHGDH-dACT mutant lost the p300/cMyc/AF9 axis (Figure 4G), which mainly explained that cMyc transactivation was impaired in the cells lossing PHGDH-ACT domain. Combined with data from the interaction between PHGDH-ACT domain and cMyc, and the transactivation of cMyc and the formation of p300/cMyc/AF9 axis regulated by PHGDH-ACT domain, all the the above results prove that PHGDH facilitates cMyc transactivation using its ACT domain to drive the formation of p300/cMyc/AF9 axis.

**Figure 5.**
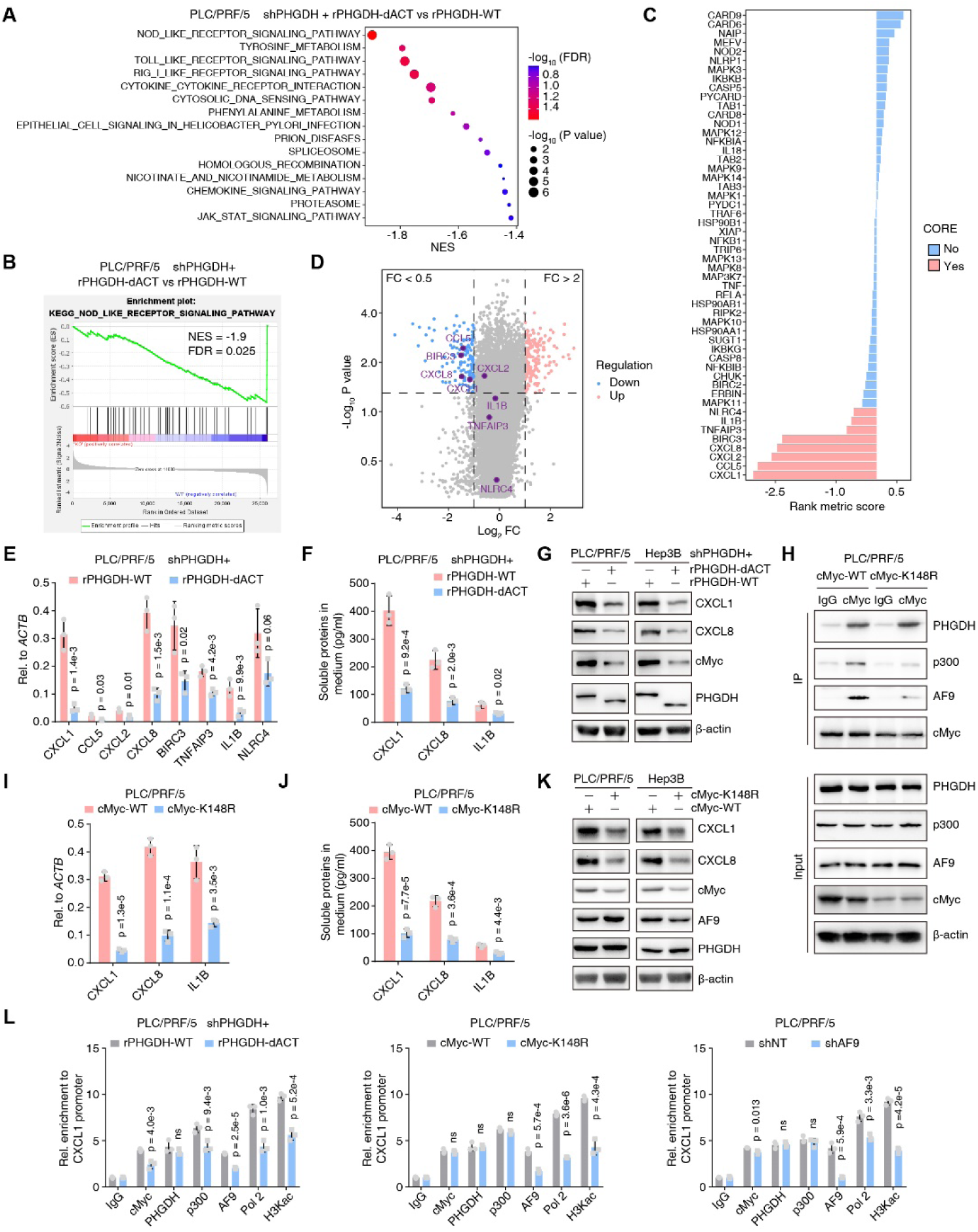
PHGDH/cMyc axis drives CXCL1/8 expression. **(A-D)** RNA-sequencing analyses were performed using PHGDH-depleted PLC/PRF/5 cells rescued with rPHGDH-WT or rPHGDH-dACT. Gene Ontology (GO) enrichment analyses of the differentially expressed genes were presented **(A)**. GSEA enrichment plot of the KEGG pathway NOD-like receptor-related gene was shown in **(B)**. The correlation of all NOD-like receptor-related gene expression with the phgdh expression status was displayed by the ranking metric score. A positive score indicates a correlation with the rPHGDH-dACT and a negative score indicates a correlation with rPHGDH-WT; The red indicates a gene that contributes most to the enrichment result and the blue indicates a gene that contributes less **(C)**. The total differentially expressed genes (FC>2 or FC<0.5; p value<0.05) were displayed using a volcano plot. FC, fold change of rPHGDH-dACT compared to rPHGDH-WT **(D). (E)** qRT-PCR validated the top up-regulated genes from **C**. (Mean ± SD, two-tailed Student’s t-test, n = 3). **(F)** ELISA examined the concentration of CXCL1/8 and IL1B in the medim culturing PHGDH-depleted PLC/PRF/5 cells, which were rescued with rPHGDH-WT or rPHGDH-dACT. (Mean ± SD, two-tailed Student’s t-test, n = 3) **(G)** Immunoblotting analysis of CXCL1/8, PHGDH and cMyc was performed using the indicated cells and antibodies. **(H)** Co-IP analysis of cMyc, p300 and PHGDH was performed using PLC/PRF/5 cells expressing WT or K148R mutant Myc. Antibody against cMyc was used to enrich cMyc associated complex. Immunoblotting analysis of PHGDH, p300, cMyc and AF9 was performed using the indicated antibodies. **(I)** qRT-PCR validated CXCL1/8 and IL1B genes using PLC/PRF/5 cells expressing WT or K148R mutant Myc. (Mean ± SD, two-tailed Student’s t-test, n = 3). **(J)** ELISA examined the concentration of CXCL1/8 and IL1B in the medim culturing PLC/PRF/5 cells expressing WT or K148R mutant Myc. (Mean ± SD, two-tailed Student’s t-test, n = 3). **(K)** Immunoblotting analysis of CXCL1/8, PHGDH and cMyc was performed using the indicated cells and antibodies. **(L)** ChIP analysis of PHGDH, cMyc, p300, RNA Pol II, AF9 and H3Kac on *CXCL1* gene promoter was performed using indicated cells. IgG was used as a blank control. (Mean ± SD, two-tailed Student’s t-test, n = 3)

### PHGDH/cMyc axis drives CXCL1/8 expression

To examine whether and how the oncogenic role of cMyc in the progression of liver cancer is regulated by PHGDH, we first analyzed the RNA profiles of PLC/PRF/5 cells expressing WT or dACT mutant PHGDH and GO enrichment analyses showed that the NOD-like receptor related pathway ranked the first in the differentially expressed genes (Figure 5A). GSEA revealed that the KEGG pathway NOD-like receptor-related genes was down-regulated in PLC/PRF/5 cells expressing dACT mutant PHGDH compared to WT (Figure 5B). The correlation of all NOD-like receptor-related gene expression with dACT mutant PHGDH was plotted by the ranking metric score. The red bars indicate genes that contribute most to the enrichment result, such as *CXCL1/2/8* and *CCL5* (Figure 5C). The total differentially expressed genes (FC > 2 or FC < 0.5; p value < 0.05) were displayed using a volcano (Figure 5D) and we marked the significant down-regulated genes, such as *CXCL1/8, CCL5* and *BIRC3*, in dACT mutant PHGDH. We tested the top 8 genes contributing most to the GSEA enrichment on the NOD-like receptor pathway by qRT-PCR. Among the 8 genes, *CXCL1/8* and *IL1B* presented the highly differential expression level (Figure 5E and Supplementary Figure 5A). We then examined the soluble levels of CXCL1/8 and IL1B in the medium culturing PLC/PRF/5 or Hep3B cells expressing WT or dACT mutant and dACT impaired the concentration of the secreted CXCL1/8 and IL1B in the medium, suggesting the decreased expression of CXCL1/8 and IL1B in liver cancer cells (Figure 5F and Supplementary Figure 5B). Immunobloting analysis further proved a dramatic reduction of CXCL1/8 after loss of PHGDH ACT domain (Figure 5G).

Our previous study reported that cMyc acetylation at K148 was required for recruitment of AF9, which is a subunit of super elongation complex and plays an aggressive role in liver cancer cells ^28^. In this study, we reinforced this idea by examining the formation of PHGDH/p300/cMyc/AF9 axis in the cells expressing WT or K148R cMyc. Of note, comparing to the interaction between PHGDH and WT cMyc, the reduced interaction between or PHGDH and K148R cMyc mainly depends on the impaired protein stability of cMyc (Figure 5H). In line with it, the cells expressing K148R cMyc mutant showed a reduced expression of *CXCL1/8* and *IL1B* (Figure 5I and Supplementary Figure 5C), also presented a reduced of secreted CXCL1/8 and IL1B in the medium (Figure 5J and Supplementary Figure 5D). Immunobloting analysis further proved the dramatic reduction of CXCL1/8 after loss of cMyc protein stability (Figure 5K).

As both AF9 and PHGDH were involved in the p300/cMyc axis, we also examined whether AF9 depletion exerted effect on *CXCL1/8* expression. As shown in Supplementary Figure 5E-F, we observed a reduction of secreted CXCL1/8 and the total CXCL1/8 protein, further supporting that there was an axis of PHGDH/p300/cMyc/AF9 as dysregulation of each one in this axis caused the same effect on CXCL1/8 expression.

Although we have identified that the subunit of PHGDH/p300/cMyc/AF9 axis has the same effect on CXCL1/8 expression, we are still not sure whether the axis directly targets *CXCL1/8* gene promoter. Then, a Ch-IP assay using antibodies separately against PHGDH/p300/cMyc/AF9, Pol 2 and H3Kac to measure the direct regulation of *CXCL1/8* gene expression. Of note, In the cells expressing PHGDH-dACT mutant, or cMyc-K148R mutant, or AF9 depletion, the association of p300/cMyc/AF9, Pol II and H3Kac with the *CXCL1/8* gene promoters was decreased in varying degrees, but the promoter of PHGDH binding target gene is not affected (Figure 5L and Supplementary Figure 5G).

Together, PHGDH/cMyc axis initiated the *CXCL1/8* gene expression by recruiting p300 and AF9, which further reinforced the expression intensity of *CXCL1/8* genes.

### Loss of nuclear PHGDH hampered TAM recruitment by liver cancer cells and impaired liver cancer progression

To test whether loss of nuclear PHGDH or PHGDH ACT domain depletion has effect on liver cancer cells, we first validated liver cancer cell proliferation using PHGDH mutant cells, rescued with PHGDH-NES or PHGDH-dACT. Compared to loss of full length of PHGDH, we observed that deletion of ACT domain slightly impaired liver cancer cell proliferation while cytosolic localized PHGDH slightly increased cell proliferation using CCK8 (Figure 6A), indicating cytosolic PHGDH facilitates cell proliferation. As liver cancer cells expressing high levels of PHGDH confer resistance to Sorafenib ^11^, we also tested whether the nuclear PHGDH contributes to Sorafenib resistance and found that under Sorafenib treatment, liver cancer cells expressing deletion of ACT domain have similar sensitivity with control cells, but the cells with cytosolic PHGDH display stronger resistance than control cells, suggesting PHGDH/cMyc axis has no effect on Sorafenib resistance (Figure 6B). Because nuclear PHGDH forms an axis with cMyc and AF9, hence, we examined Sorafenib resistance in the cells expressing WT or K148R cMyc, or AF9 depletion. As shown in Supplementary Figure 6A and B, we observed no obvious difference on Sorafenib resistance in the above groups, which were in line with PHGDH mutants.

**Figure 6.**
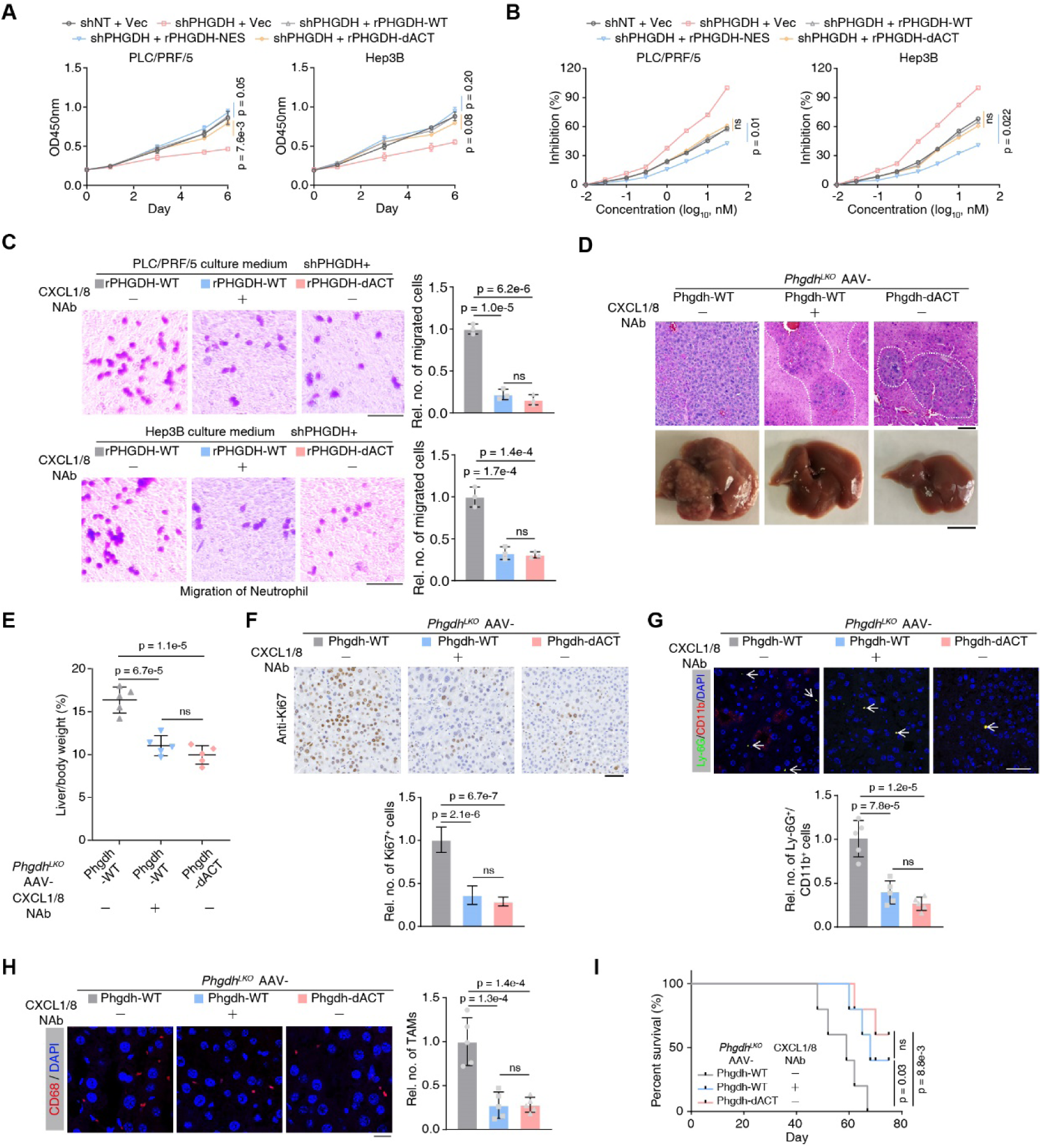
Destruction of PHGDH/cMyc axis hampered neutrophil recruitment by liver cancer cells and then impaired liver cancer progression. **(A)** Cell proliferation was evaluated by spectrofluorometer at wavelength OD450 in PHGDH-depleted PLC/PRF/5 or Hep3B cells rescued with Vec, rPHGDH-WT, rPHGDH-NES and rPHGDH-dACT. Cells expressing shNT rescued with Vec were used as control. (Mean ± SD, two-way ANOVA). **(B)** Sorafenib inhibition was evaluated by trypan blue staining in PHGDH-depleted PLC/PRF/5 or Hep3B cells rescued with Vec, rPHGDH-WT, rPHGDH-NES and rPHGDH-dACT. Cells expressing shNT rescued with Vec were used as control. (Mean ± SD, two-way ANOVA). **(C)** Neutrophil recruitment was evaluated by cell migration, which was performed by placing the medium from culturing PHGDH-depleted PLC/PRF/5 or Hep3B cells in the lower well and neutrophil cells in the upper transwell chamber. Scale bars: 20 μm. (Mean ± SD, one-way ANOVA followed by Tukey’s multiple comparisons test, n = 3 per group). **(D)** Macroscopic images and H&E staining of *Phgdh*^*LKO*^ mouse livers under different AAV treatments (Phgdh-WT with or without neutrolized antibodies against CXCL1/8 and Phgdh-dACT) at W9 after hydrodynamic injection of MET/CAT constructs and the SB transposase. Scale bars: 100 μm (upper); 1 cm (lower). **(E)** Liver/body weight ratios of MET/CAT-transfected mice under different AAV treatments (Phgdh-WT with or without neutrolized antibodies against CXCL1/8 and Phgdh-dACT) at W9 were measured. (Mean ± SD, one-way ANOVA followed by Tukey’s multiple comparisons test, n = 5 per group). **(F-H)** IHC or IF staining of Ki67, Ly6G/CD11b and CD68 in mouse liver sections at W9 after *Phgdh*^*LKO*^ mice receving different AAV treatments (Phgdh-WT with or without neutrolized antibodies against CXCL1/8 and Phgdh-dACT). The relative number of Ki67^+^ cells **(F)**, neutrophil **(G)** or TAMs **(H)** was calculated. Scale bars: 100 μm. **(F)**, 20 μm **(G-H)**. (Mean ± SD, one-way ANOVA followed by Tukey’s multiple comparisons test, n = 5 per group) **(I)** Kaplan-Meier plot showing the survival of MET/CAT-transfected mice under different AAV treatments (Phgdh-WT with or without neutrolized antibodies against CXCL1/8 and Phgdh-dACT). (Log-rank test, n = 5 per group).

In addition to hepatocytes, there are non-parenchymal cells in liver tissue, especially macrophages and neutrophils, which form a immune-network to mediate immune cell infiltration in tumor stroma and play as a vital role in driving tumor progression ^16,17^. Additionally, CXCL1/8, secreted by macrophages, monocytes, or other cells direct the recruitment of neutrophils in the hepar and promotes liver cancer progression ^29–33^. Compared to control liver cancer cells, the cells with antibody neutralizing treatment or genetic manipulation of PHGDH-dACT reduce the recruitment of the neutrophils (Figure 6C). We also obtained the consistent result from the cells expressing K148R cMyc, or AF9 depletion (Supplementary Figure 6C and D).

To further verify whether the destruction of PHGDH/cMyc axis hampers liver cancer progression, we delivered AAV carrying the Phgdh-WT or the Phgdh-dACT mutant under the regulation of the thyroid hormone binding globulin promoter into *Phgdh*^*LKO*^ adult mice. These mice were subjected to the MET/CAT to induce hepatocarcinogenesis and neutralizing antibodies (NAbs) against CXCL1/8 were applied to treat above mice. As expected, the Phgdh-WT group with NAb treatment and the Phgdh-dACT mutant group evidently reduced the formation of tumor nodules when compared to the Phgdh-WT group without NAb treatment (Figure 6D). Consistently, mice from both the Phgdh-WT group with NAb treatment and the Phgdh-dACT mutant group showed lower liver/body ratios (Figure 6E).

The staining of Ki67, Ly6G/CD11b and CD68 respectively shows that, in the same region of liver tumor, the proliferation rate of liver cancer cells, the number of neutrophils and TAM recruitment were decreased when PHGDH/cMyc axis was destroyed (Figure 6F, G and H). Mice from both the Phgdh-WT group with NAb treatment and the Phgdh-dACT mutant group showed increased survival when compared to the Phgdh-WT group without NAb treatment (Figure 6I). Data above proves that PHGDH/cMyc axis in the liver cancer cells is required for recruiting neutrophils and TAM, which further facilitate liver cancer progression.

### Nuclear PHGDH predicts poor prognosis and associates with cMyc in clinic

To discover the clinic significance of PHGDH, we mined PHGDH expression in liver tumor tissue and normal adjacent tissues (NATs). As shown in Figure 7, A and B, the RNA and protein levels of PHGDH were downregulated in NATs (Figure 7, A and B). Generally, the downregulation of gene expression indicates this gene would play a suppressive role in tumor development. However, both this study and the previous study ^11^ confirmed that PHGDH contributes to liver cancer progression. We then performed IHC using 25 paired liver tumor samples and found a consistent result that the total PHGDH was downregulated in the tumor tissues but its major location has changed from cytosol to nucleus (Figure 7C).

**Figure 7.**
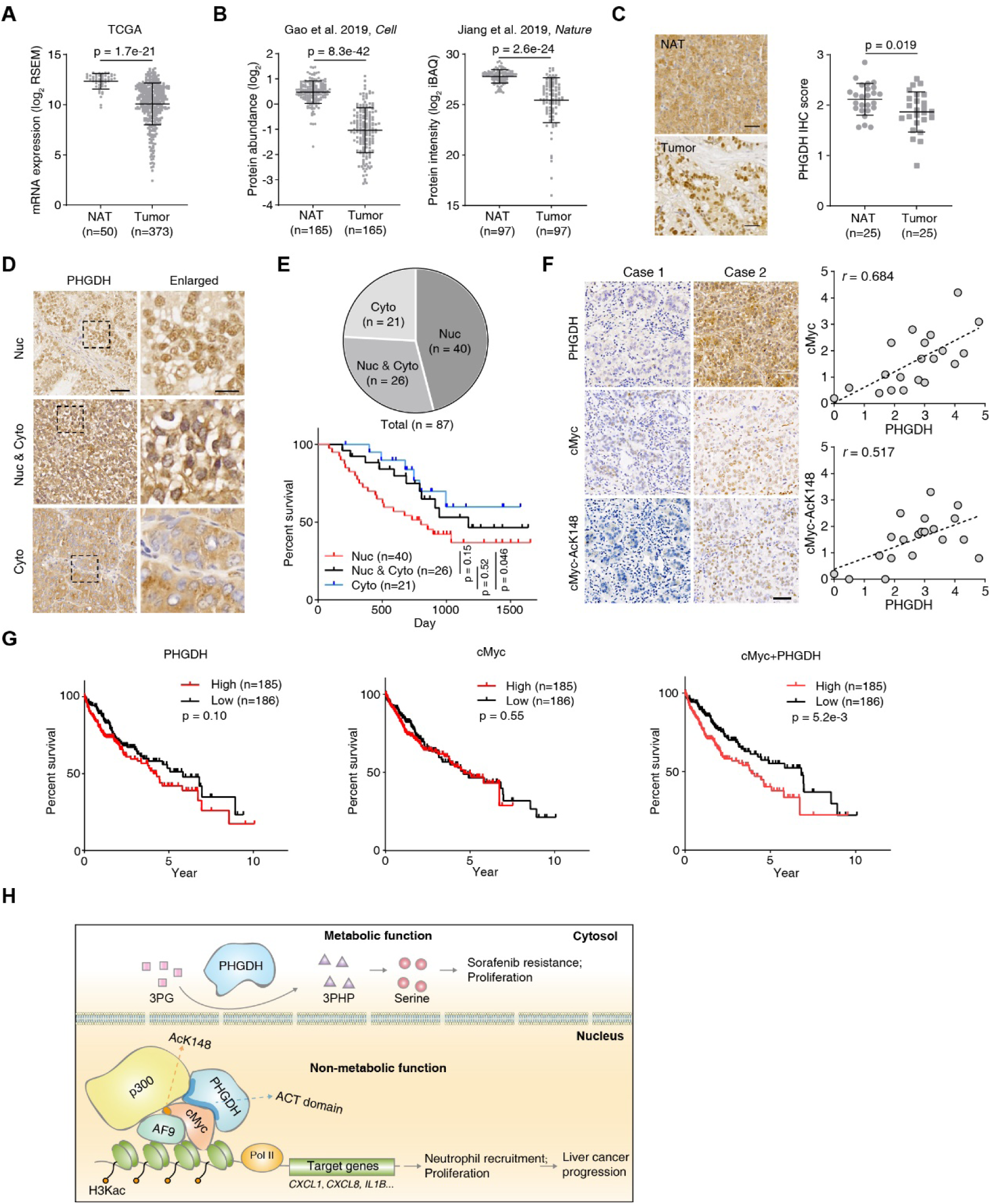
Nucleus-localized PHGDH predicts poor prognosis and associates with cMyc in clinic. **(A)** Comparation of *PHGDH* expression levels in HCC tumors and normal adjacent tissues (NATs) using the TCGA dataset. (Mann-Whitney test) **(B)** Comparation of PHGDH protein levels in HCC tumors and paired normal adjacent tissues (NATs) using the two HCC proteomic datasets (Gao et al. 2019, *Cell*; Jiang et al. 2019, *Nature*). (Wilcoxon matched-pairs signed rank test) **(C)** IHC staining of PHGDH in paired human liver tissues (NAT) and liver cancer tissues (n = 25, Wilcoxon matched-pairs signed rank test). Representative images of PHGDH staining in paired samples were shown. Scale bars: 50 μm. **(D)** IHC staining of PHGDH in human liver cancer tissues. Representative images of nucleus (Nuc), cytoplasma (Cyto), both nucleus and cytoplasma (Nuc & Cyto) localized PHGDH were shown. Scale bars: 50 μm (left); 10 μm (right). **(E)** The patient counts of different subcellular localized PHGDH in human liver cancer tissues (upper). Kaplan-Meier plot showing the survival of patients with different subcellular signals of PHGDH. (Log-rank test) (lower). **(F)** The positive correlation of PHGDH and cMyc, or PHGDH and cMyc-AcK148 in clinical liver cancer patients were examined by IHC. Right panels show the semi-quantitative scoring (using a scale from 0 to 5) between two staining signals was carried out (Pearson product moment correlation test). Scale bar, 50 μm **(G)** Survival plot of PHGDH expression, cMyc expression, the combined expression of PHGDH and cMyc in TCGA-LIHC data with median cutoff. (Log-rank test.) **(H)** Work model of the nonmetabolic role of PHGDH.

To further validate the localization of PHGDH in human liver cancer tissues, we analyzed 87 tumor samples that showed positive signals. Based on the subcellular localization of the IHC signals of PHGDH, these tissues can be categized into three types: mainly nucleus localized (Nuc), both nucleus and cytoplasm localized (Nuc & Cyto) and mainly cytoplasm localized (Cyto) (Figure 7D). Notably, about half of the samples showed shown strong nucleus-localized PHGDH signals (Figure 7E, upper panel). More importantly, we found that patients with strong nucleus-localized PHGDH showed worse associated with a poor prognosis when compared to patients with cytoplasm-localized PHGDH (Figure 7E, lower panel).

In order to verify whether PHGDH and cMyc have synergistic effects in clinic, we first performed IHC with clinical liver cancer samples using antibodies against PHGDH, cMyc and cMyc-AcK148. We observed a positive correlation between PHGDH and cMyc, or PHGDH and cMyc-AcK148, suggesting PHGDH has a regulatory role on cMyc protein stability in clinical (Figure 7F). We also mined the TCGA data and evaluated the association of the expressions of PHGDH, cMyc and PHGDH/cMyc with patients’ survival respectively and found that the combined expression of PHGDH and cMyc has significance to predict liver cancer patients’ prognosis (Figure 7G). Thus, these results indicate that the translocation of PHGDH form cytoplasm to nucleus is an unfavorable biomarker for liver cancer patients. These findings further suggest that PHGDH and cMyc exert synergistic effects on liver cancer progression in clinic.

## Discussion

In this study, we discovered a distinct function of PHGDH which used its ACT domain to promote cMyc transactivation on the promoter of genes required for recruiting neutrophils and supporting TAM filtration in the liver cancer tissue, and then drove liver cancer into adcanced stages (Figure 7H).

Integrative transcriptomic and proteomic analyses have been shown to be valuable tools for investigating biological issues ^34,35^. We combined these two omics methods to examine the molecular events underlying MET/CAT-induced liver carcinogenesis and ultimately identified the SSP through dynamic proteome profiling, which improved the reliability of our study.

As summarized previously ^6^, hydrodynamic transfection of MET/CAT with SB is a reliable and widely used method to generate liver cancer in mice. Recently, this method has been used to facilitate understanding of hepatocarcinogenesis in many works ^36–38^. Here, we found that the Afp, Glul and cytochrome C family genes were gradually up-regulated in a mouse model during liver cancer progression (Figure 1, B and C). More importantly, two of the three SSP enzymes, Phgdh and Psat1, showed robust upregulation. In view of the importance of the SSP, in which Phgdh is the first rate-limiting enzyme, we specifically deleted an exon of *Phgdh* in mouse livers. We performed tissue-specific deletion because systemic knockout of *Phgdh* is lethal ^9^. To ensure the reliability of our MET/CAT-induced liver cancer model, we tested the activity of AST and ALT in the livers of *Phgdh*^*LKO*^ mice and found no significant differences in these mice compared to the control mice, indicating that loss of Phgdh does not cause abnormal liver function in the absence of MET/CAT.

Given the oncogenic role of Phgdh, Phgdh inhibitors have been designed to hamper various diseases. A few inhibitors of PHGDH are currently available, such as CBR-5884 ^39^, NCT-502 and NCT-503 ^23^. In this study, the enzyme activity of Phgdh was suppressed by NCT-503 or manupilated by AAV introduced enzyme-dead PHGDH in mouse livers, but this inhibition did not evidently slow the progression of liver cancer. Therefore, we speculated that Phgdh has a nonmetabolic function in driving liver cancer progression.

We explored the interaction networks of Phgdh to uncover its nonmetabolic role and found that Phgdh associated with cMyc during different stages of MET/CAT-driven liver carcinogenesis. A few previous reports have demonstrated that metabolic enzymes can regulate the transactivation of transcription factors. For example, FBP1 interacts with and inhibits HIF1α ^15^, while PKM2 interacts directly with the HIF1α and promotes transactivation of HIF1α target genes ^40^. The identification of cMyc as an interactor of PHGDH is a key step in this study. cMyc directly regulates more than one thousand target genes that affect many aspects of tumor behavior, such as proliferation, growth, metastasis, metabolic abnormality and drug resistance ^41^.

The nucleus colocalization of PHGDH and cMyc is the basis for their interaction. We further detected the localization of PHGDH in clinical human liver cancer samples, and found that 66 out of 87 samples showed nucleus PHGDH signals. More significantly, the nucleus localization of PHGDH is correlated with worse prognosis, suggesting that PHGDH functionally contributes to tumor progression in a location dependent manner. Metabolic enzymes enter the nucleus to participate in regulating gene expression or mRNA stability. Our previous study showed that UGDH localizes in the nucleus and binds to HuR, eliminating UDP-glucose-mediated inhibition of HuR ^42^.

PHGDH facilitated cMyc expanding its scope of targeting genes, such as CXCL1/2/5/8, IL1B and CCL5. All of these chemokines presented the capcity of recruiting immune cells in the TME. Chemokines can indirectly modulate tumor growth through their effects on tumor stromal cells and immune cells by inducing the release of growth and angiogenic factors from cells in the TME. In the TME, tumor cells or other cells have been shown to acquire the ability to produce growth-promoting chemokines which forms networks with chemokine receptors. Understanding the interplay between chemokine-chemokine receptor networks between the tumor cells and their microenvironment is a novel approach to overcome the problem of metastatic heterogeneity. Recent advances in the understanding of chemokine networks pave the way for developing a potential targeted therapeutic strategy to treat advanced cancer. Previous studies have identified the hepatic expressed CXCL1/5 contribute to neutrophil infiltration in the liver, which promotes steatosis-to-NASH progression in HFD-fed mice ^43^ or advanced stages of liver cancer ^44^. Blockage of these chemokines-chemokine receptor networks significantly improved the treatment of the liver diseases ^45,46^. Additionally, Mollaoglu et al. use genetically engineered mouse models of non-small-cell lung cancer (NSCLC) to reveal that the neutrophil recruitment determines the nature of the tumor ^47^, suggesting the classification of liver cancer stages could be also achieved by examining the infiltration of neutrophils in liver.

It is worth noting that some chemokines, such as CXCL8, have a dual role in promoting or suppressing cancer. During the progression of the tumor, elevated levels of CXCL8 have been detected in tumors and potential function as a mitogenic and angiogenic factor ^48,49^. While dying cancer cells can be immunogenic and can direct the antitumor immune response. CXCL8, for example, can increase the immunogenicity of dying cancer cells by translocating calreticulin to the cell surface ^50^.

Although cMyc controls many aspects of tumorigenic processes, the efforts to use cMyc as a therapeutic target have been quite frustrating, indicating there are still some unsolved mysteries to understand cMyc transactivation ^51–53^. Our study provides a perspective to inhibit cMyc transactivation by disrupting PHGDH/cMyc axis in liver cancer cells. What is more reliable is that when we integrately analyzed the expression levels of PHGDH and cMyc, we find that the two genes have a high clinical synergy and can predict the survival of liver cancer patients.

In line with the evidence that PHGDH localized in nucleus at the cellular level, we also obseverd the nuclear location of PHGDH in the both human and mouse advanced liver cancer. The nuclar intensity of PHGDH could mark the progression of liver cancer and PHGDH/cMyc axis shows a new layer of metabolic enzymes in regulating the reciprocal action of tumor cells and microenvironment.

## Methods

### Antibodies and reagents

Rabbit polyclonal or monoclonal antibodies against PHGDH (PA5-82863, 1:50 for ChIP), PSAT1 (PA5-22124, 1:200 for IHC), CD68 (14-0688-82, 1:500 for IHC), Ki67 (MA5-14520, 1:500 for IHC), Lamin B (33-2000, 1:1,000 for IB), and mouse monoclonal antibody against PHGDH (MA5-31357, 1:200 for IHC, 1:1,000 for IB, 1:100 for IP), CXCL1 (MA5-23811, 1:1000 for IB, 1:200 for ELISA), CXCL8 (M801, 1:1000 for IB, 1:200 for ELISA), Ly-6G (14-5931-82, 1:100 for IF), CD11b (53-0112-82, 1:100 for IF) and CD68 (MA5-13324, 1:100 for IF) were purchased from Thermo Fisher. The mouse monoclonal antibody against FLAG (F3165, 1:5,000 for IB, 1:100 for IP) and the rabbit polyclonal antibody against AF9 (HPA001824, 1:1000 for IB, 1:100 for ChIP) were purchased from Sigma-Aldrich. Antibody against HA (#3724, 1:2,000 for IB) was purchased from Cell Signaling Technology (CST). Mouse monoclonal antibodies against β-actin (60008-1-Ig, 1:5,000 for IB), GST tag (66001-2-Ig, 1:5,000 for IB, 1:100 for IP) and His tag (66005-1, 1:3,000 for IB) was purchased from Proteintech. Mouse monoclonal antibodies against cMyc (sc-40, 1:2,000 for IB, 1:200 for IHC and 1:50 for ChIP), tubulin (sc-5286, 1:2,000 for IB) and CYP2B10 (sc-73546, 1:50 for IHC) were purchased from Santa Cruz Biotech. A histone H3Kac antibody (Clone 2G1F9, 1:200 for ChIP, a POL2 antibody (Clone 4H8, 1:100 for ChIP) and a p300 antibody (Clone NM11, 1:200 for ChIP, 1:1000 for IB, 1:100 for IP) were purchased from Active Motif. The antibody against cMyc-AcK148 was prepared in a previous study ^28^. NCT-503 (SML1659) and NCT-503 Inactive Control (SML1671) were purchased from Sigma-Aldrich. siRNAs targeting human EP300 (106443) was purchased from Thermofisher.

### Experimental mice

The *Phgdh*^*LKO*^ (*Phgdh*^*fl/fl*^:*Alb-Cre*) mouse line was generated on a mixed FVB/N and C57BL/6 background by breeding *Phgdh*^*fl/fl*^ mice with *Albumin-Cre* transgenic mice. C57BL/6 mice were used for MET/CAT model construction in the proteomic study. All animal studies were conducted on male mice aged 6–24 weeks. The mice were group-housed (4–5 mice per cage) except for fewer than 5% of the mice, which were single-housed later because of the death of cage mates. All mice were maintained under a 12-hour light/dark cycle with free access to water and standard mouse chow. All mice received humane care, and all experimental procedures were approved by the Institutional Animal Care and Use Committee (IACUC). For hydrodynamic injection, plasmids (Met: PT3EF1aH-hMet; Ctnnb1/b-catenin: PT3EF1aH-b-catenin; SB transposase: pCMV/SB) were kindly provided by Professor Lijian Hui (Shanghai Institute of Biochemistry and Cell Biology, Chinese Academy of Sciences). Oncogene-expressing constructs were delivered by hydrodynamic tail vein injection into male mice at 6 weeks of age, as described previously ^20,54–57^. Antibodies against CXCL1/8 were injected into vein with a concentration of 1 μg/50 μl, once every two days for two weeks to neutralize the soluble CXCL1/8 in the liver.

### Human samples

Human samples were collected at The First Affiliated Hospital, Sun Yat-sen University. The HCC samples were collected during surgery. Written informed consent was obtained from all patients or their guardians for the use of the biospecimens for research purposes, which were carried out in accordance with the approved guidelines “Use of experimental animals and human subjects”. All procedures were approved by the institutional review board of Zhongshan School of Medicine, Sun Yat-sen University.

### Cell lines

PLC/PRF/5 and Hep3B cells were obtained from the American Type Culture Collection (ATCC, USA). All cells were authenticated using the short tandem repeat (STR) method and tested negative for mycoplasma.

### Total RNA isolation, RNA-seq and data analysis

Total RNA was extracted from the samples using Trizol Reagent (Invitrogen). Purified RNAs were treated with RNase R (Epicenter, 40 U, 37°C, 3 h), followed by purification with Trizol. Subsequently, using the NEBNext UltraTM RNA Library Prep Kit, RNA-seq libraries were prepared and subjected to deep sequencing with an Illumina HiSeq 3000 at Seqhealth Technology Co., Ltd., Wuhan, China. Paired-end 150 bp read length sequencing was performed. Trimmed reads were aligned to the mouse genome (Mm10, Genome Reference Consortium GRCm38) using TopHat v2.0.6 ^58^. FPKM (fragments per kilobase of exon per million fragments mapped) values were calculated by Cufflinks using default parameters for gene expression levels. Differential expression was defined using the indicated fold-changes and false discovery rate (FDR) 0.05. The RNA-seq raw data, related to Figure 1 and Figure 5, have been deposited to Gene Expression Omnibus (GEO) (GEOAccession numbers: pending, should be available in 5-6 working days).

### Protein extraction and peptide digestion

The animal studies were performed in compliance with the regulations and guidelines of Guangzhou University. Liver tissues were quickly collected by multispot sampling and pooled after MET/CAT model mice were sacrificed at W0, W2 and W7. The proteins were extracted with SDS lysis buffer (100 mM dithiothreitol, 4% sodium SDS, 100 mM Tris-HCl, pH 7.6) using mechanical homogenization followed by ultrasonication. The proteins were then denatured and reduced at 95°C for 5 min. The insoluble debris was removed by centrifugation at 12,000 × *g* for 10 min, and the supernatant was used for proteomic experiments. The protein concentration was determined by a tryptophan-based fluorescence quantification method. The filter-aided sample preparation (FASP) method was used for protein digestion as previously described ^59^. Briefly, 100 μg of protein was loaded into a 10 kDa centrifugal filter tube (Millipore), washed twice with 200 μl of UA buffer (8 M urea in 0.1 M Tris-HCl, pH 8.5), alkylated with 50 mM iodoacetamide in 200 μl of UA buffer for 30 min in the dark, washed thrice with 100 μl of UA buffer again and finally washed thrice with 100 μl of 50 mM NH_4_HCO_3_. After each of the above steps, the samples were centrifuged at 12,000 × *g* at 25°C. The proteins were digested with trypsin (Promega Corporation, USA) at an enzyme-to-substrate ratio of 1:50 (w/w) in 200 μl of 50 mM NH_4_HCO_3_ at 37°C for 16 hours, and the peptides were eluted by centrifugation. The digested peptides were desalted using C18 Stage Tips and evaporated to dryness in a Speed-Vac sample concentrator. Finally, the amounts of purified peptides were determined with a NanoDrop (Thermofisher, USA).

### TMT labeling and high-pH RP fractionation

An isobaric labeling experiment was conducted according to the TMT kit instructions. Each channel was labeled with 20 μg peptides. The three samples for W0 were labeled with channels 126, 128N and 129C; those for W2 were labeled with 127N, 128C and 130N; and those for W7 were labeled with 127C, 129N and 130C. Channel 131 was used to label a sample of mixed peptides from W0, W2 and W7. The TMT reagents (0.8 mg) were dissolved in anhydrous acetonitrile (41 μl) and added to the peptides (dissolved in 100 μl of 100 mM TEAB). The labeling reactions were incubated for 1 hour at room temperature, and then 8 μl of 5% hydroxylamine was added to the samples for 15 min to quench the reaction. The labeled peptides were pooled, vacuum-centrifuged to dryness and subjected to C18 solid-phase extraction desalting (3M Empore). High-pH reversed-phase (RP) LC was used for peptide fractionation. The labeled peptides were fractionated using a Waters XBridge BEH300 C18 column (150×1 mm, OD 5 μm) at a flow rate of 0.2 ml/min on an Agilent 1200 LC instrument. The mobile phase contained solvent A (10 mM NH_4_COOH, adjusted to pH 10.0 with NH_3_·H_2_O) and solvent B (90% acetonitrile [ACN], 10 mM NH_4_COOH, adjusted to pH 10.0 with NH_3_·H_2_O). A 110-min gradient was set as follows: 1%–5% B in 2 min, 5%–25% B in 35 min, 25%–40% B in 43 min, 40%–55% B in 6 min, 55%–95% B in 3 min, 95% B for 4 min, 95%–1% B in 1 min, and 1% B for 16 min. The eluate was collected every 2 min, and the eluate samples were combined by a concatenation strategy into 25 fractions. The fractions were vacuum-centrifuged to dryness and then subjected to MS analysis.

### LC-MS/MS data acquisition

LC-MS/MS analysis was carried out with an EASY-nLC 1000 liquid chromatograph (Thermo Fisher Scientific) coupled to an Orbitrap Fusion mass spectrometer (Thermo Fisher Scientific). For each fraction, peptides (∼1 μg) were loaded onto an in-house packed analytical column (75 μm ID × 20 cm, ReproSil-Pur C18-Pur, 3 μm, Dr. Maisch GmbH, Ammerbuch, Germany) and subjected to a 120-min gradient at a flow rate of 300 nl/min. The column was heated to 50°C using a column compartment to prevent overpressure during LC separation. Mobile phase A consisted of 0.1% formic acid, and mobile phase B consisted of 0.1% formic acid in 100% ACN. The LC gradient was set as follows: 4%–28% B in 95 min, 28%– 40% B in 15 min, 40%–100% B in 2 min, and 100% B for 8 min. The spray voltage was set at 2,500 V in positive ion mode, and the ion transfer tube temperature was set at 275°C. Data-dependent acquisition was performed using Xcalibur software with the profile spectrum data type. The MS1 full-scan parameters were set to a resolution of 120,000 at m/z 200, a lens radio frequency (RF) of 60%, an automatic gain control (AGC) target of 4e5 and maximum injection time (IT) of 50 ms by Orbitrap mass analyzer (350–1700 m/z). ‘Top-speed’ MS2 scans were generated by higher-energy C-trap dissociation (HCD) fragmentation at a resolution of 50,000 at m/z 200 with an AGC target of 1e5 and a maximum IT 100 ms. The function “inject ions for all available parallelizable time” was enabled. The isolation window was set at 1.2 m/z. The normalized collision energy (NCE) was set at 38%, and the dynamic exclusion time was 40 s. Precursors with charges of 2–6 were included for MS2 analysis.

### Proteomic database searching

All mass spectrometric data were analyzed using MaxQuant 1.6.1.0 against the mouse Swiss-Prot database containing 22,259 sequences (downloaded in August 2019). TMT-MS2 was chosen for proteomic quantification with a reporter ion mass tolerance of 0.003 Da. Carbamidomethyl cysteine was searched as a fixed modification. Oxidized methionine and protein N-term acetylation were set as the variable modifications. The enzyme specificity was set to trypsin/P. The maximum missed cleavage sites was set to 2. The tolerances of the first search and the main search for peptides were set to 20 ppm and 4.5 ppm, respectively. The minimal peptide length was set to 7. The false discovery rate (FDR) cutoff for peptides and proteins was set to 0.01. The mass spectrometry proteomics data have been deposited to the ProteomeXchange Consortium via the PRIDE partner repository with the dataset identifier PXD017809.

### Proteomic data analysis

The Persus, Excel and R software programs were used for MS data analysis. The correlation matrix, PCA plot, HCA plot and cluster profile were created in Persus. The significance of the biological process enrichment analysis results for the clusters in Figure 1D were calculated by Fisher’s exact test in Persus. The KEGG pathway and GO biological process enrichment analyses were performed on the Database for Annotation, Visualization and Integrated Discovery (DAVID) website, and the adjusted p-values from Benjamini-Hochberg correction are reported. The protein-protein interaction maps were drawn from the STRING database. The volcano plot was generated in Excel.

### qRT-PCR

Total RNA was prepared from cell samples using TRIzol (Invitrogen, USA) according to the manufacturer’s protocol. Reverse transcriptase PCR was performed using M-MLV reverse transcriptase (Promega, USA). qRT-PCR was performed using SYBR^®^ Premix Ex Taq™ (Takara, Japan) and 300 nmol/L of each primer. Amplification was performed with a 7500 Fast Real-Time PCR Systems (Applied Biosystems, USA) according to the manufacturer’s protocol. The data were normalized to the expression of the control gene (β-actin) for each experiment. The data are presented as the mean ± SD from three independent experiments. The sequences of the primer pairs used for qRT-PCR are listed in Supplemental Table 2.

### Histology and immunohistochemistry

Mouse liver samples and human liver cancer samples were fixed overnight in 4% PFA (4°C) and embedded in paraffin blocks. Immunohistochemistry staining and hematoxylin and eosin staining were performed as previously described ^60,61^. Tissue sections were stained with the indicated antibodies. We quantitatively scored the tissue sections according to the percentages of positive cells and the staining intensity and determined the localization of the signals as described previously ^60,61^.

### Measuring reactive oxygen species (ROS) in liver tissue

The ROS level of liver tissue were measured using 2,7-dichlrofluorescein diacetate (DCFDA), which is converted to highly fluorescent DCF by cellular peroxides (including hydrogen peroxide). The assay was performed as described before ^62,63^. In brief, 1% tissue homogenates were prepared in ice-cold 40 mM Tris–HCl buffer (pH 7.4). Samples were further diluted to 0.25 % with the same buffer and placed on ice. Each sample was divided into two equal fractions (2 ml each). To one fraction, 40 μl of 1.25 mM DCFDA in methanol was added for ROS measurement. The same volume of methanol was added to the other fraction as a control for sample autofluorescence. All samples were incubated for 15 min in a 37 °C water bath. The fluorescence was determined at 488 nm excitation and 525 nm emission using a Bio-Rad fluorometer.

### Measuring 3PG, 3PHP and Ser in liver tissue

Tissue samples were prepared as previously described ^64^. 3PG, 3PHP and serine levels were determined by standard ion-exchange chromatography using a Beckman 6300 automated amino acid analyzer.

### Immunoprecipitation, immunoblotting analysis and LC-MS

Immunoprecipitation was performed with lysates from the indicated cultured cells and was followed by IB with corresponding antibodies. Briefly, after trypsinization, cells were harvested and washed twice with cold PBS. The cell pellets were resuspended and lysed in lysis buffer (Millipore). Then, the lysates were sonicated with a Covaris S220 to destroy chromatin and centrifuged at 15,000 × *g* to remove the cell debris. The protein concentration was determined using a BCA Protein Assay Kit (Pierce) according to the manufacturer’s instructions. A total of 100 mg of protein was incubated with the indicated antibodies overnight and then mixed with protein A or protein G-agarose beads (Millipore), respectively. The immunocomplexes were collected by centrifugation at 1,000 × *g*, resolved by SDS–PAGE and subsequently transferred to PVDF membranes (Millipore). The blots were blocked with 5% nonfat milk and then incubated with primary antibodies and HRP-conjugated secondary antibodies. The blots were developed using SuperSignal West Pico (Thermo Scientific) and detected with a Tanon 6200 Luminescent Imaging Workstation. For LC-MS of the immunocomplexes, the proteins were resolved by SDS-PAGE and subjected for in-gel digestion by trypsin as the widely used protocol ^65^. The resulted peptides were separated by a LC gradient of two hours and analyzed on a Q Exactive HF mass spectrometer (Thermo Fisher Scientific). The MS data were searched against the mouse database using the MaxQuant software. The iBAQ (intensity-based absolute-protein-quantification) intensity was used for comparison between samples.

### DNA constructs and stable cell lines

PCR-amplified PHGDH and cMyc were cloned into pCDH, pCDH-3’Flag or 3’HA. The target sequence of PHGDH is listed in Supplemental Table 1, as is a nontargeting sequence used as a negative control. The cMyc reporter plasmid was the same as those used in a previous study ^66^. Cells cultured in Dulbecco’s modified Eagle’s medium (DMEM) or Roswell Park Memorial Institute (RPMI) 1640 medium were incubated with a retrovirus encoding shRNA or pCDH for 8 hours. Forty-eight hours after infection, the cells were selected by culture with 2 mg/ml puromycin.

### Cell culture and transfection

Cells were cultured as described previously ^60^. Above cells were maintained in DMEM or RPMI medium supplemented with 10% fetal bovine serum (FBS). Cell transfection was performed with Lipofectamine™ 3000 (Invitrogen, USA) according to the manufacturer’s instructions.

### Subcellular fractionation analyses

PLC/PRF/5 and Hep3B cells were collected and washed three times with cold PBS. Nuclear or cytosolic fractions were prepared using a Nuclear Extract Kit (Active Motif, USA).

### Immunofluorescence staining

Cells were fixed and incubated with anti-PHGDH and anti-cMyc antibodies, Alexa Fluor dye-conjugated secondary antibodies and DAPI according to standard protocols. Liver tissues were fixed in 4% formaldehyde and embedded in optimum cutting temperature (OCT) compound (Thermo) after dehydration by 30% sucrose solution. The tissue sections were were washed in PBS, 0.3% Triton X-100 and blocked in PBS with 10% normal goat serum followed by incubation with anti-Ly-6G, anti-CD11b or anti-CD68 antibodies. After washing, samples were incubated with Alexa Fluor dye-conjugated secondary antibodies and DAPI. The imaging was performed on an Olympus FV3000 confocal laser scanning microscope (Olympus).

### GST pull-down assay

Purified proteins, GST-ACT (1 μg) and His-cMyc (1 μg) were incubated at room temperature for 2 hours, and then GST beads were added to pull down the GST-tagged proteins for an additional 2 hours. The GST beads were washed with lysis buffer (Millipore) three times for 5 min each time. Then, the GST beads bound with proteins were boiled for 8 min after adding SDS loading buffer.

### Luciferase reporter gene assay

The transcriptional activation of cMyc in liver cancer cells was measured using a Dual-Luciferase Assay Kit (Promega) on a GloMax 20/20 luminometer (Promega, E1910) following the manufacturer’s instructions. In detail, the cells were plated in triplicate at a density of 2 × 10^4^ cells/well in a volume of 500 μl in separate 24-well microtiter plates. After transfection with the indicated plasmids for 48 hours, the cells were washed with cold PBS and lysed with lysis buffer. The relative levels of luciferase activity were normalized to the levels in untreated cells and to the levels of Renilla luciferase activity of the control plasmid in each group.

### ChIP assay

For ChIP experiments, cells were fixed in 1% formaldehyde for 10 min for pull-down of PHGDH, RNA Pol II, AF9, cMyc, p300 and H3Kac. The cells were rotated in cold lysis buffer (10 mM Tris-HCl pH 7.4, 10 mM NaCl, 3 mM MgCl_2_, 0.5% NP40, supplemented with freshly prepared PMSF and protease inhibitor cocktail) for 30 min. Then, the nuclear pellets were resuspended in RIPA buffer (300 mM NaCl, 3 mM EDTA, 1% NP40, 0.5% sodium deoxycholate, 0.1% SDS, 100 μg/ml BSA, 50 mM Tris-HCl pH 7.5) and sonicated with a Covaris S220 to yield DNA fragments of approximately 200–500 bp. A ChIP-grade antibody was incubated with 30 μl of protein G Dynabeads (Thermo Fisher) for 4 hours. Then, the DNA fragments were coimmunoprecipitated with the specific antibody-conjugated protein G beads at 4°C overnight. The DNA was purified with a MinElute PCR Purification Kit (Qiagen, Germany). The promoter regions of target genes were amplified and quantified by qRT-PCR using SYBR Green (Takara, Japan) on an ABI 7500 Fast system. For PCR, 0.02 ng of the immunoprecipitated DNA and 2 ng of the total DNA were used in a 20 μl reaction. The results from each immunoprecipitation were normalized to the respective inputs. The primers are listed in Supplemental Table 2.

### Cell proliferation assay

Cell proliferation was analyzed with CCK-8 assay. Cells were seeded into 96-well plates (500 cells/well), then incubated at 37 °C for the next 6 days. CCK-8 assay was performed conforming to the manufacturer’s instructions every two days. The OD value for each well was measured at 450 nm with a reference at 650 nm using a microtiter plate reader (Becton Dickinson).

### Sorafenib sensitivity assays

To determine the sensitivity of liver cancer cells to Sorafenib, 2×10^4^ of live tumor cells were seeded in 96 well plates and treated with different concentrations of Sorafenib (from 0 to 100 nM). After 72 hours, the cell viability was determined by Trypan blue staining assay and inhibition efficency was calculated using Graphpad Prism 7.

### Neutrophil transwell assays

Neutrophils were isolated according to the methods described previously ^67^ and cultured with or without CXCL1/8 neutrolizing antibodies (1:50 by volume). Transwell assays were performed on 24-well plate with inserts (BD Biosciences) according to manufacturer’s instruction. Briefly, 5×10^4^ of neutrophils were cultured in the upper chamber and allowed to migrate for 36 hours before fixation for crystal purple staining. The lower culture media was obtained by culturing PLC/RF/5 and Hep3B in DMEM media containing 10% FBS for 36 hours and then used for the cell migration transwell assay.

### PHGDH expression analysis of the previous published clinical datasets

The TCGA data (Figure 7A) was downloaded from https://gdac.broadinstitute.org/. The protein data (Figure 7B) were from two proteomic datasets of HCC, Gao et al. 2019, *Cell* ^68^, and Jiang et al. 2019, *Nature* ^69^.

### Survival analysis and gene expression correlation analysis

Survival analysis was performed in GraphPad Prism software, and the log-rank test p-values are reported. TCGA-LIHC gene expression and survival data for PHGDH and cMyc were downloaded from www.tcgaportal.org. Survival plots for PHGDH and cMyc was reconstructed in GraphPad with median cutoffs. To create Figure 7G, we performed a combined survival analysis of PHGDH and cMyc. Given the different expression levels of the two proteins, we ranked the abundances of PHGDH and cMyc and then summed the ranks as the combined abundance (PHGDH+cMyc) levels to enable comparison.

### AAV virus in vivo delivery

For AAV virus infection, 2 × 10^11^ genomic particles of AAV-Vec, AAV-Phgdh-WT, AAV-Phgdh-ED AAV-Phgdh-dACT were reconstituted in 200 μl PBS and injected intravenously through tail veil injection with BD Ultra-Fine Insulin Syringes.

### Statistical analysis

All data are presented as the mean ± SD unless stated otherwise. Statistical calculations were performed using GraphPad Prism software (version 8.0.2). Unpaired Student’s t-test and one-way ANOVA followed by multiple comparison test (Dunnett’s or Tukey’s method) were used to calculate statistical probability in this study. Two-tailed log-rank test was used to analyze survival data. The number of animals used for each experiment is indicated in the figure legends.

## Author contributions

XW, WZ, and HZhou designed this study. HZhu and HY performed most of experiments and analyzed the data. YC helped with some expriments. XW, HZhu, and WZ wrote and revised the manuscript. All authors approved the final manuscript.

## Acknowledgements

This study is supported by the National Science Fundation of China (No.92053113) and the Hundred Talent Program of Guangzhou University, the National Key Research and Development Program from the Ministry of Science and Technology of China (No.2017YFC1700200) and the Yangfan Project of Shanghai Science and Technology Commission (No.20YF1457300).

## Notes

**Conflict of interest:** The authors have declared that no conflict of interest exists.

### Competing Interest Statement

The authors have declared no competing interest.

