## Supplementary material for "Nuclear PHGDH promotes neutrophil recruitment to drive liver cancer progression": Sup information

### Supplemental Figure Legends

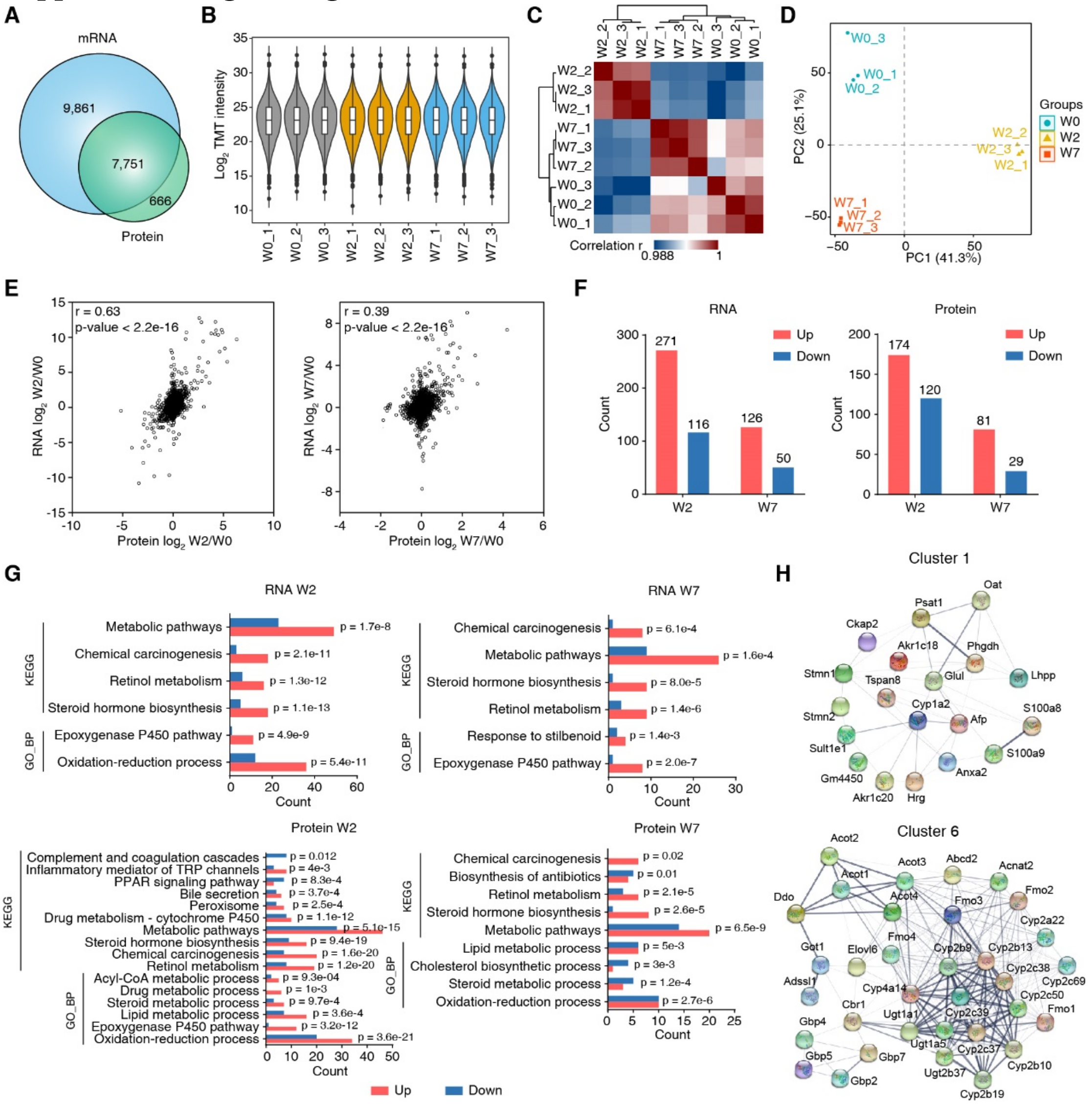


**Fig. S1. Overall quality and uniformity of the proteomic and RNA profile data.**

(*A*) Overlap of the identified mRNA and proteins at the gene level. A total of 17,612 RNAs and 8,417 proteins were quantified at all three time points.

(*B*) Boxplot of the log2 transformed TMT intensity of the proteomic data.

(*C*) Correlation matrix of the proteomic data. The three repeats for the same time point were clustered into a subgroup.

(*D*) Principal component analysis of the proteomic data. The three repeats for the same time point were clustered together.

(*E*) Correlations between RNA and protein levels at W2 (C) and W7 (D) compared to W0. The Pearson correlation coefficient and p-value were calculated.

(*F*) Counts of up- and down-regulated genes and proteins. Red indicates upregulation, and blue indicates downregulation.

(*G*) KEGG pathway and GO biological process enrichment analysis results for RNA sequences and proteins regulated in W2 vs. W0 and in W7 vs. W0. The x-axis represents the gene count. The enrichment analysis was performed in the DAVID, and the adjusted p-value of each item is labeled on the right side.

(*H*) Protein-protein interaction maps of the proteins in Cluster 1 and Cluster 6 were drawn using the STRING database. The line thickness between every two proteins indicates the strength of the interaction.

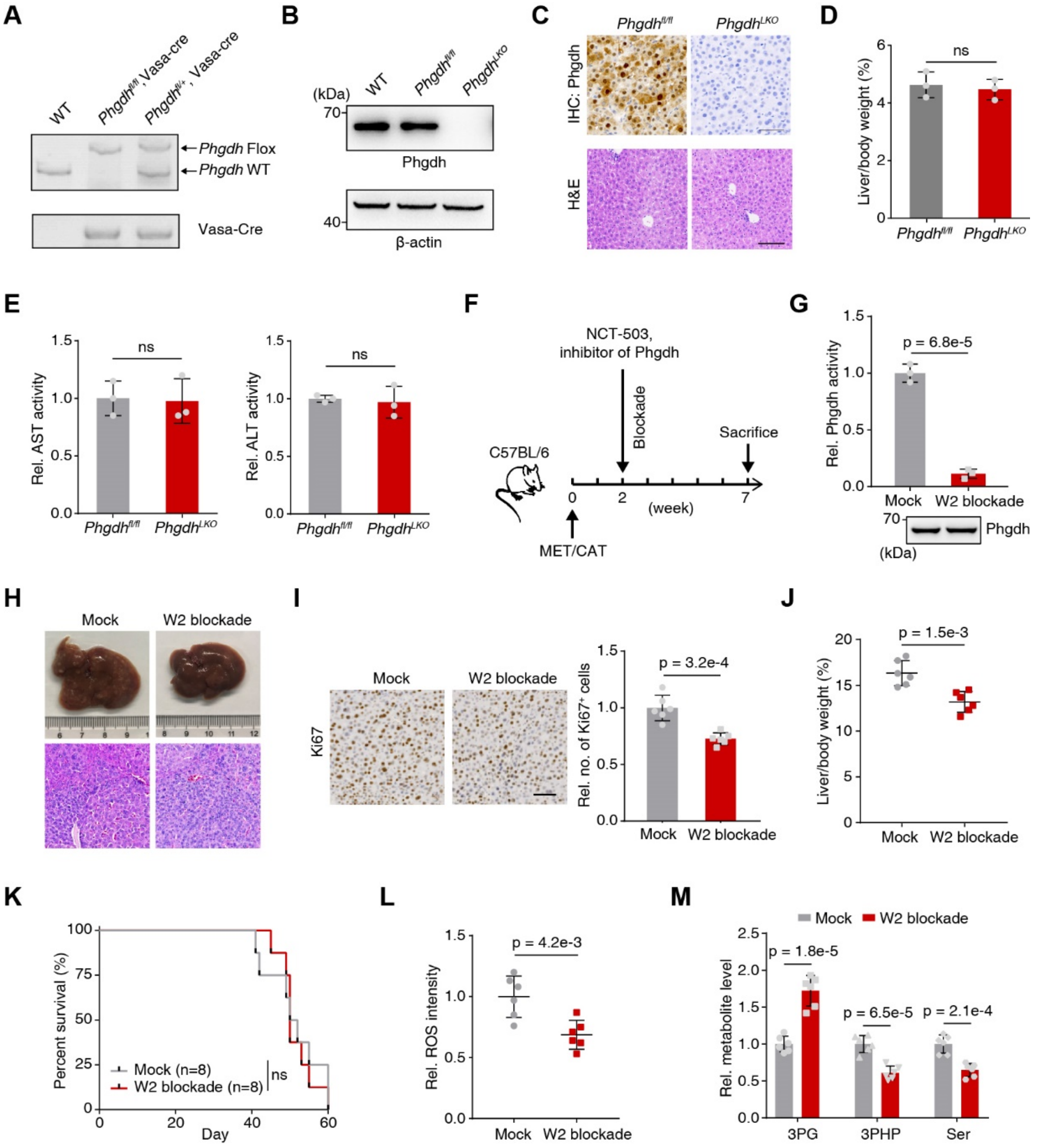


**Fig. S2. Hepatic loss of *Phgdh* enhances mouse survival after MET/CAT-driven**

**hepatocarcinogenesis.**

(*A*) Genotypes were determined by PCR using liver genomic DNA.

(*B*) Phgdh protein in liver tissues of *Phgdh*$^{fl/fl}$ and *Phgdh*$^{LKO}$ mice was detected by immunoblotting. Loading control, β–actin.

(*C*) IHC staining of Phgdh and H&E staining using liver sections from the indicated mice at 6-week-old.

(*D*) The liver/body weight ratios of the indicated mice at 6-week-old were measured. (Mean ± SD, two-tailed Student's t-test, n = 3; ns, not significant.)

(*E*) The serum ALT and AST levels were measured in the indicated mice at 6-week-old without MET/CAT induction. (Mean ± SD, two-tailed Student's t-test, n = 3)

(*F*) Schematic diagram of NCT-503 treatment in mice with MET/CAT-driven liver cancer. Mice received NCT-503 treatment at W2. The mock group received saline at W2 and served as a negative control. The mice were sacrificed at W7. Dosage, 100 mg/kg.

(*G*) Phgdh was immuno-precipitated from livers and incubated with its substrate 3PG and $NAD^+$ for testing the enzymatic activity. The protein levels were adjusted to the same extent and confirmed by immunoblotting analysis. (Mean ± SD, two-tailed Student's t-test, n = 3)

(*H*) Macroscopic images and H&E staining of MET/CAT-transfected mouse (Mock vs. W2 blockade) liver sections at W7.

(*I*) Ki67 staining of MET/CAT-transfected liver sections under mock or W2 blockade treatments (left panel). Relative numbers of Ki67-positive cells in liver sections (right panel) were shown. Scale bars, 200 μm. (Mean ± SD, two-tailed Student's t-test, n = 6)

(*J*) The liver/body weight ratios of MET/CAT-transfected mice (Mock vs. W2 blockade) were measured at W7. (Mean ± SD, two-tailed Student's t-test, n = 6.)

(*K*) Kaplan-Meier plot showing the survival of MET/CAT-transfected mice (Mock vs. W2 blockade) over 120 days. (Log-rank test.)

(*L*) The relative ROS intensities were measured in MET/CAT-transfected liver samples (Mock vs. W2 blockade) at W7. (Mean ± SD, two-tailed Student's t-test, n = 6.)

(*M*) The relative levels of three metabolites (3PG, 3PHP and Ser) were measured in

MET/CAT-transfected liver samples (Mock vs. W2 blockade) at W7. (Mean ± SD, two-tailed Student’s t-test, n = 6.)

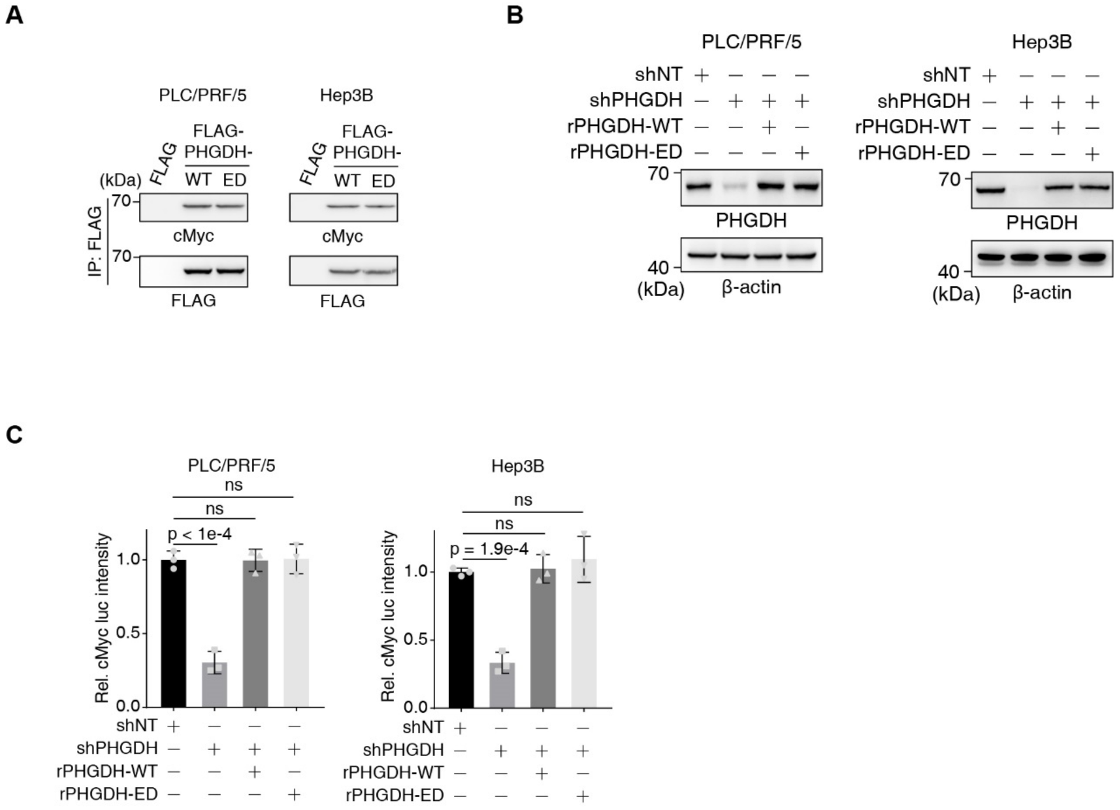


**Fig. S3. Related to Fig. 3.**

(*A*) Co-IP was performed with an antibody against FLAG in PLC/PRF/5 and Hep3B (FLAG tagged PHGDH-WT and -ED) cells. An antibody against cMyc was used to detect the association between FLAG-PHGDH and cMyc.

(*B*) PLC/PRF/5 or Hep3B cells stably expressing shNT or shPHGDH were rescued with rPHGDH-WT or rPHGDH-ED.

(*C*) cMyc transactivation in PHGDH-depleted PLC/PRF/5 or Hep3B cells rescued with rPHGDH-WT or rPHGDH-ED was measured with a Dual-Luciferase® Reporter Assay System according to the manual.

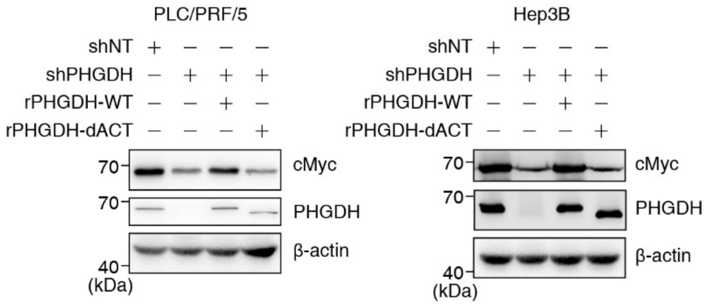


**Fig. S4. Related to Fig. 4.** (*A*) PLC/PRF/5 or Hep3B cells stably expressing shNT or shPHGDH were rescued with rPHGDH-WT or rPHGDH-dACT.

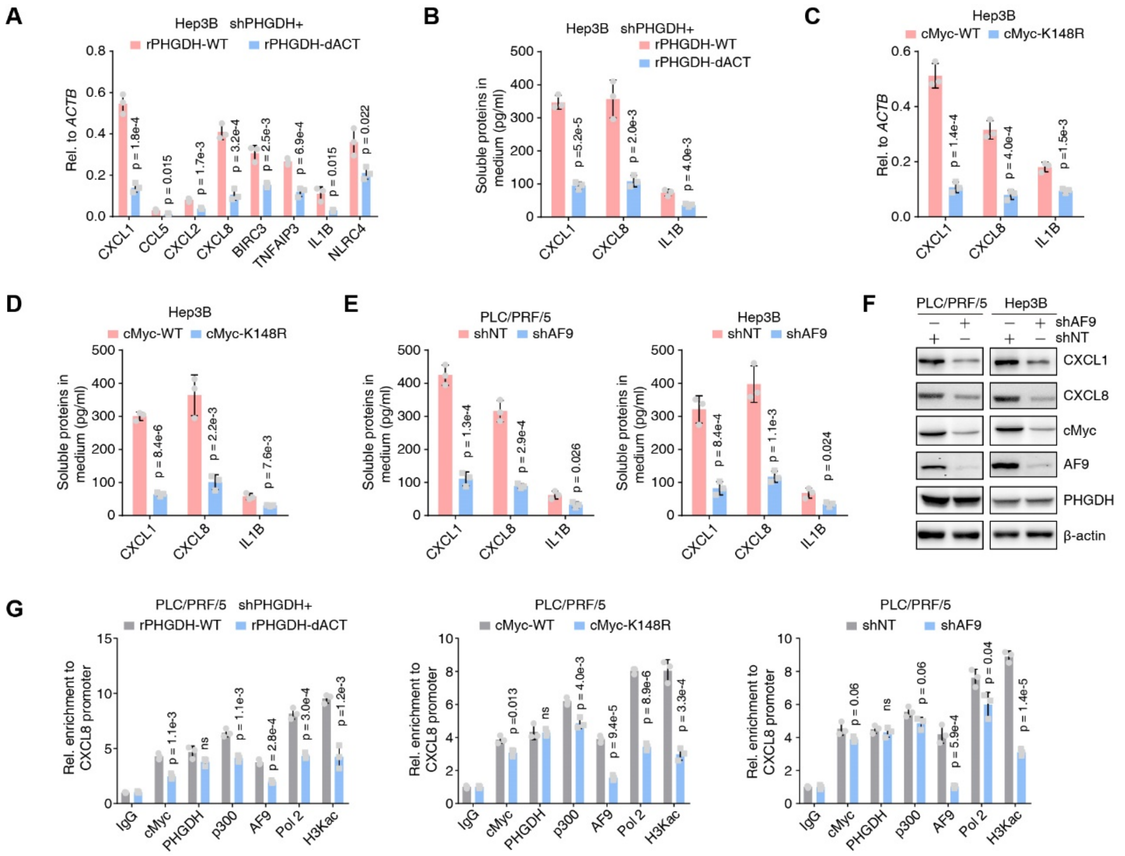


**Fig. S5. Related to Fig. 5.**

(*A*) qRT-PCR validated the top up-regulated genes from Figure 5C using PHGDH-depleted Hep3B cells, which were rescued with rPHGDH-WT or rPHGDH-dACT. (Mean ± SD, two-tailed Student's t-test, n = 3).

(*B*) ELISA examined the concentration of CXCL1/8 and IL1B in the medium culturing PHGDH-depleted Hep3B cells, which were rescued with rPHGDH-WT or rPHGDH-dACT.

(*C*) qRT-PCR validated CXCL1/8 and IL1B genes using Hep3B cells expressing WT or K148R mutant Myc. (Mean ± SD, two-tailed Student's t-test, n = 3).

(*D*) ELISA examined the concentration of CXCL1/8 and IL1B in the medium culturing Hep3B cells expressing WT or K148R mutant Myc. (Mean ± SD, two-tailed Student's t-test, n = 3).

(*E*) ELISA examined the concentration of CXCL1/8 and IL1B in the medium culturing PLC/PRF/5 or Hep3B cells expressing shNT or shAF9. (Mean ± SD, two-tailed Student's t-test, n = 3).

(*F*) Immunoblotting analysis of CXCL1/8, PHGDH, AF9 and cMyc was performed using the indicated cells and antibodies.

(*G*) ChIP analysis of PHGDH, cMyc, p300, RNA Pol II, AF9 and H3Kac on CXCL1 gene promoter was performed using indicated cells. IgG was used as a blank control. (Mean ± SD, two-tailed Student's t-test, n = 3).

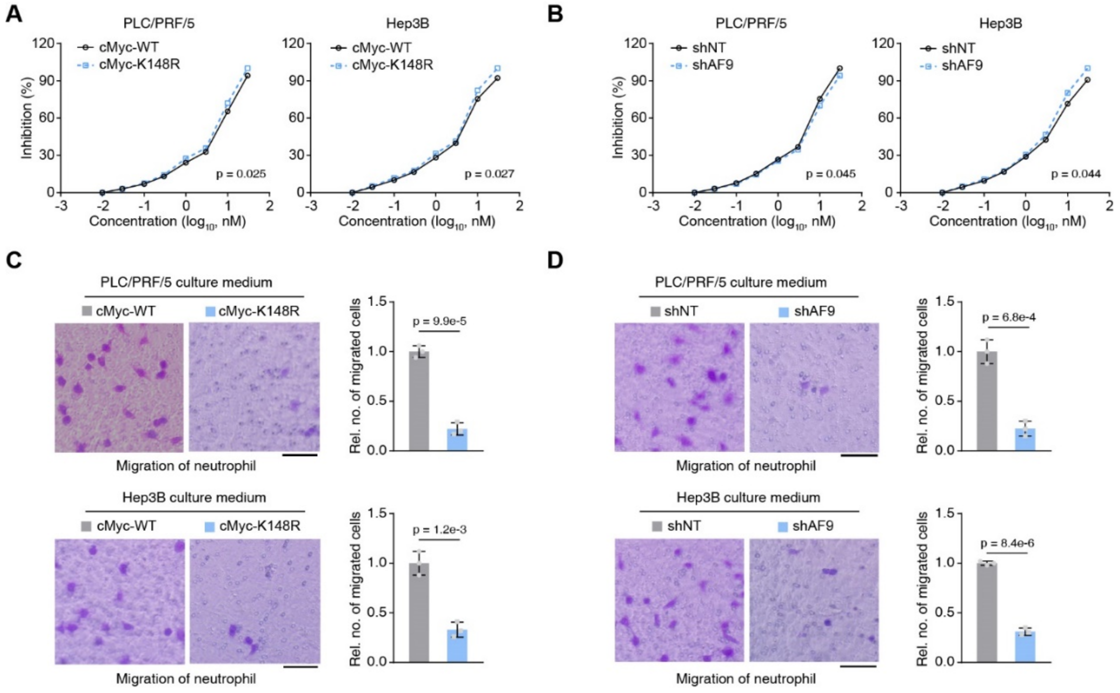


**Fig. S6. Related to Fig.6.**

(*A*) Sorafenib inhibition was evaluated by trypan blue staining in PLC/PRF/5 or Hep3B cells expressing WT or K148R mutant cMyc. (two-way ANOVA).

(*B*) Sorafenib inhibition was evaluated by trypan blue staining in PLC/PRF/5 or Hep3B cells expressing shNT or shAF9. (two-way ANOVA).

(*C*) Neutrophil recruitment was evaluated by cell migration， which was performed by placing the medium from culturing PLC/PRF/5 or Hep3B cells expressing WT or K148R mutant cMyc in the lower well and neutrophil cells in the upper transwell chamber. Scale bars: 20 µm. (Mean ± SD, two-tailed Student's t-test, n = 3).

(*D*) Neutrophil recruitment was evaluated by cell migration， which was performed by placing the medium from culturing PLC/PRF/5 or Hep3B cells expressing shNT or shAF9 in the lower well and neutrophil cells in the upper transwell chamber. Scale bars: 20 µm. (Mean ± SD, two-tailed Student's t-test, n = 3).

**Supplemental Tables**

**Supplemental Table 1. The sequence (5' -> 3') used for silencing genes:**

shNT : targeting sequence, CAACAAGATGAAGAGCACCAA

shPHGDH (TRCN0000028548):

CCGGAGGTGATAACACAGGGAACATCTCGAGATGTTCCCTGTGTTATCACCTTTTTT

**Supplemental Table 2. The list of primers used in this study.**

1. The sequence (5' -> 3') of primers used for qRT-PCR:

Primers for human genes:

CXCL1-forward: GCAGACCCTGCAGGGAATTCA

CXCL1-reverse: GCTTTCCGCCCATTCTTGAGT

CCL5-forward: CCAGCAGTCGTCTTTGTCAC

CCL5- reverse: CTCTGGGTTGGCACACACTT

CXCL2-forward: GCAATCCCCGGCTCCTGCGG

CXCL2-reverse: GGTGAATTCCCTGCAGGGTC

CXCL8-forward: GTGTGTAAACATGACTTCCA

CXCL8- reverse: GCACTGACATCTAAGTTCTTTA

BIRC3-forward: AAGCTACCTCTCAGCCTACTTT

BIRC3-reverse: CCACTGTTTTCTGTACCCGGA

TNFAIP3-forward: TTGTCCTCAGTTTCGGGAGAT

TNFAIP3- reverse: ACTTCTCGACACCAGTTGAGTT

IL1B-forward: AGCTACGAATCTCCGACCAC

IL1B- reverse: CGTTATCCCATGTGTCGAAGAA

NLRC4-forward: TGCATCATTGAAGGGGAATCTG

NLRC4-reverse: GATTGTGCCAGGTATATCCAGG

Primers for mouse genes:

Phgdh-forward: CCTCATTGTCCGGTCTGCTAC

Phgdh-reverse: CATCTTTCATCGAAGCTGTTGC

Psat1-forward: CAGTGGAGCGCCAGAATAGAA

Psat1-reverse: CCTGTGCCCCTTCAAGGAG

S100a8-forward: AAATCACCATGCCCTCTACAAG

S100a8-reverse: CCCACTTTTATCACCATCGCAA

S100a9-forward: GCACAGTTGGCAACCTTTATG

S100a9-reverse: TGATTGTCCTGGTTTGTGTCC

Cyp2b9-forward: GCTCATTCTCTGGTCAGATGTTT

Cyp2b9-reverse: CGCTTGTGGTCTCAGTTCCA

Cyp2b10-forward: TGCTGTCGTTGAGCCAACC

Cyp2b10- reverse: CCACTAAACATTGGGCTTCCT

Cyp2b13-forward: AGTGAGCCACGAGACTTCATC

Cyp2b13-reverse: GTGCTCCAGTAGGGACAATATCT

Cyp2b19-forward: TTCCAGTGCATCGTAGCAAAT

Cyp2b19- reverse: GGAACTCTGTATGGTGGTTGG

Cyp2c37-forward: GCCAATCCTTCACCAATTTA

Cyp2c37- reverse: GCCTCGTGTTTTTCCATACA

Cyp2c38-forward: CACGGCCCATTGTTGTATTGC

Cyp2c38-reverse: TGAGTGTGAAACGTCTTGTCTCT

Cyp2c39-forward: GAGGAAGCATTCCAATGGTAGAA

Cyp2c39- reverse: TGTGAAGCGCCTAATCTCTTTC

Cyp2c50-forward: ACTGTGGTGTTGCATGGATATG

Cyp2c50- reverse: GAGAAGCGCCTTGTGTTTTTC

Cyp4a14-forward: TCTGGGTTCTTCCAATGGGC

Cyp4a14-reverse: GGACTCGTATATTGCTCCCCG

2. The sequence (5' -> 3') of primers used for Chip-qPCR:

Human:

CXCL1-forward: GTTCTCAGGGATCCGCCCCA

CXCL1-reverse: GGAGAGAGCAGCGCGGGCCA

CXCL8-forward: GATTGGCTGGCTTATCTTCA

CXCL8- reverse: GTGCTCCGGTGGCTTTTTATA
